# G4-Attention: Deep Learning Model with Attention for predicting DNA G-Quadruplexes

**DOI:** 10.1101/2024.11.04.621797

**Authors:** Shrimon Mukherjee, Pulakesh Pramanik, Partha Basuchowdhuri, Santanu Bhattacharya

## Abstract

G-quadruplexes (G4s) are the four-stranded non-canonical nucleic acid secondary structures, formed by the stacking arrangement of the guanine tetramers. They are involved in a wide range of biological roles because of their exceptionally unique and distinct structural characteristics. After the completion of the human genome sequencing project, a lot of bio-informatics algorithms were introduced to predict the active G4s regions *in vitro* based on the canonical G4 sequence elements, G-*richness*, and G-*skewness*, as well as the non-canonical sequence features. Recently, sequencing techniques like G4-seq and G4-ChIP-seq were developed to map the G4s *in vitro*, and *in vivo* respectively at a few hundred base resolution. Subsequently, several machine learning and deep learning approaches were developed for predicting the G4 regions using the existing databases. However, their prediction models were simplistic, and the prediction accuracy was notably poor. In response, here, we propose a novel convolutional neural network with Bi-LSTM and attention layers, named G4-Attention, to predict the G4 forming sequences with improved accuracy. G4-Attention achieves high accuracy and attains state-of-the-art results in the G4 propensity and mismatch score prediction task in comparison to other available benchmark models in the literature. Besides the balanced dataset, the developed model can predict the G4 regions accurately in the highly class-imbalanced datasets. Furthermore, the model achieves a significant improvement in the cell-type-specific G4 prediction task. In addition, G4-Attention trained on the human genome dataset can be applied to any non-human genomic DNA sequences to predict the G4 formation propensities accurately. We have also added interpretability analysis of our model to gain further insights.

**Author summary:** G-quadruplex, a non-canonical secondary nucleic acid structure, has emerged as a potential pharmacological target because of its significant implication in several human diseases including cancer, aging, neurological disorders, etc. Despite numerous computational algorithm developments, the prediction of G4 regions accurately in different organisms including humans still remains a challenging task. To address this, in this work, we have presented a novel advanced deep learning architecture called G4-Attention for predicting DNA G-quadruplexes in different organisms including humans. To the best of our knowledge, we are the first to incorporate Bi-LSTM and attention layers on top of a CNN architecture in a deep learning model (G4-Attention) for predicting G4-forming sequences. Our developed model outperforms existing algorithms and achieves current state-of-the-art (SOTA) results in G4 propensity and mismatch score prediction tasks. In addition, the developed model achieves superior results across non-human genomes, class-imbalanced datasets, and cell line-specific datasets. Lastly, G4-Attention can identify key features for understanding the G4 formation mechanism.

## Introduction

Guanine-rich DNA and RNA sequences have the ability to fold into four-stranded non-canonical secondary structures known as G-quadruplexes (G4s) [1]. These functional secondary structures were discovered in the late 80’s [2] and have displayed important cellular roles in living cells [3]. G4s are structurally a self-stacked arrangement of the G-quartets, a square planar organized structure formed by four guanine nucleotides connected through Hoogsteen-type of hydrogen bonding, one on top of another via *π*-*π* stacking interaction (Fig. 1) [4, 5]. These structures can form kinetically but may be stabilized thermodynamically in the presence of monovalent cations such as Na^+^, and K^+^ under physiological conditions [6, 7]. Even though G4s are spontaneously formed from four consecutive tracts of three or more guanines separated by loops having various lengths under *in vitro* conditions, many non-canonical G4s, that is G4s with longer loops, mismatches, and bulges, have been recently reported. It indicates that sequence patterns are less strict characteristics for the formation of G4s than previously believed [8, 9]. Depending upon the physiological conditions, syn- or anti-conformation of the glycosidic bond, and intrinsic features of the sequence, G4s can adopt various topologies such as parallel, anti-parallel, and hybrid G4s [10]. In addition, G4s can be formed from one (unimolecular), two (bimolecular), or four (tetramolecular) distinct strands with different topologies [11, 12]. Furthermore, the structural integrity is affected by the number of consecutive G-quartets, loop length, and the composition of nucleotides in loop regions [13, 14].

**Fig 1.**
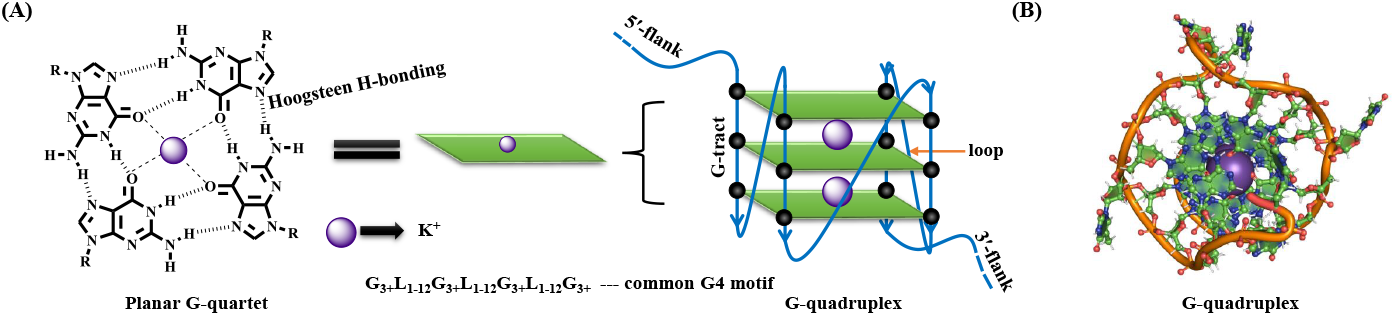
Schematic representation of canonical G-quadruplex structures in DNA. (A) Structure of planar G-quartet formed by four guanine bases through Hoogsteen type of H-bonding, and a central cation usually K^+^. The G-quadruplex structures are commonly comprised of three stacks planar G-quartet top to one another via *π*− *π* stacking interaction. (B) Top-view of human promoter G4 found in c-MYC gene (PDB : 1XAV).

The G4 motifs were significantly distributed in the telomeric regions of chromosomes, gene promoter regions, DNA replication origins, 5’-untranslated region (UTR), first intron, and exon regions of various genes [15–19]. Recently, G4s were visualized using an immunofluorescence assay, which demonstrates their existence in cells and tissues [20–22]. Several investigations have further revealed their potential involvement in critical biological functions such as telomeric stability, DNA replication, gene transcription, mRNA translation, and DNA repair [23–28]. For example, in 85-90% of cancer cells, up-regulation of telomerase enzyme was observed which blocks the activation of cell apoptosis machinery by restoring the telomeric length during the cell division process [29, 30]. The terminal 3’-telomeric DNA can down-regulate the telomerase activity by forming G4s; thereby uncontrolled tumor cell proliferation is inhibited by the activation of apoptosis machinery [29]. Furthermore, G4s can repress the expression of various oncogenes such as MYC, KIT, Bcl-2, KRAS, HRAS, and VEGF found in cancer cells [31–36]. G4s are therefore believed to be associated with human diseases like cancer, neurological disorders, and aging and have emerged as a potential target for therapeutic intervention [27, 37–39].

Because of the significant regulatory roles of G4s in cellular processes, numerous mathematical and computational algorithms were developed to predict the location of active putative quadruplex sequence (PQS) throughout the human genome with high accuracy [40]. The first PQS detection tool, quadparser was developed using the classical regular expression matching technique in C++ and Python scripting language [41]. This tool was used to detect all matching occurrences of canonical G4 motifs across the human genome by using a strict sequence pattern *G*_3−5_*N*_1−7_*G*_3−5_*N*_1−7_*G*_3−5_*N*_1−7_*G*_3−5_; where G and N represent guanine, and any other nucleotide respectively. This model identified around 376,000 unimolecular PQS motifs in the human genome (*hg19* reference). One of the main flaws of this tool is that it provides a yes/no binary output without considering the structural stability and *in-vivo* folding propensity. Since 2005, methods that were developed on the PQS motifs, which fit the regular expression mentioned above, have been used in the majority of the research studies to predict the G4 propensity [42–44]. More recently, a similar kind of regular expression matching tool, AllQuads has been developed to identify the inter-molecular G4s [45]. Meanwhile, several in-vitro and in-vivo experiments proved the existence of non-canonical i.e., imperfect G4s along with canonical G4s in the human genome [46]. Afterward, new methods were developed to identify imperfect G4s by incorporating the features of the newly found imperfect G4 and by updating the algorithms incrementally. For instance, ImGQfinder [47] can detect the non-canonical intra-molecular PQS motif with a single bulge or mismatch, and pqsfinder [48, 49] recognizes G4s folded from G-runs with flaws such as bulges, mismatches, and G4s with longer loops. In addition, pqsfinder [48, 49] also gives each hit an integer score, which represents the predicted stability of the folded G4s. Further, an algorithm based on a sliding window approach, G4Hunter [50–52] was developed to compute the quadruplex propensity score based on G-*richness* and G-*skewness* of a given sequence. Overall, these tools contributed significantly to the G4 exploration. However, due to the lack of vast biophysical and biochemical databases, these tools cannot perfectly predict the G4s sequences and stability. Recently, G4-seq [53] and G4-ChIP-seq [54] types of experimental methods were implemented to map G4s *in vitro* and *in vivo* respectively. Following that, new machine and deep learning types of artificial intelligence approaches were used to predict the G4s much more accurately using the sequencing databases [55, 56]. For example, Quadron [57], a machine learning algorithm, was trained using a large database of experimental G4 regions, obtained by the G4-seq techniques in the human genome. This algorithm can detect the G4 sequences and assign a score of the detected G4s, which represents their formation propensity *in vitro*. Other deep learning models, such as PENGUINN [58], and G4Detector [59], a convolutional neural network (CNN) were also developed to predict the G4 regions in vitro. Since all of these models were constructed throughout the training process with the G4-seq database, new models were then going to be developed for predicting G4 sequences in cells using G4-ChIP-seq peak DNA sequences. For instance, DeepG4 [60] used a CNN to predict cell-type specific active G4 regions, which is trained using a combination of G4-seq, and G4 ChIP-seq DNA sequences with chromatin accessibility measures (ATAC-seq). Further, epiG4NN [61] was developed which uses a ResNet [62] based architecture for G4s prediction in cells. Despite the advancement of numerous machine and deep learning models using the existing databases, the prediction accuracy was quite poor and requires further improvement. In addition, all the prediction models were very simplistic. The prediction accuracy can be enhanced by incorporating additional layers, such as GRU, LSTM, attention, and dilated convolution layers [63], into the networks. Developing more sophisticated models is imperative for achieving improved prediction accuracy.

In this study, we proposed a novel architecture named G4-Attention to predict the G4-forming sequences with improved prediction accuracy. Our work makes the following contributions:

- To the best of our knowledge, we are the first ones to introduce Bi-LSTM and attention layers on top of CNN to predict the G4-forming sequences. We apply our model G4-Attention to various types of problems related to learning of G4 forming sequence, like G4 propensity prediction or G4-mismatch score prediction tasks.
- G4-Attention achieves current state-of-the-art (SOTA) results in the G4 propensity prediction and G4 mismatch score prediction tasks. Particularly for the G4 propensity prediction task, G4-Attention produces an average improvement of 3.26% and 3.99% on two stabilizers *K*^+^ and *K*^+^ + PDS respectively against the second best model G4Detector. Also for G4 mismatch score prediction, G4-Attention improves second best model, G4mismatch results by 3.57% and 3.30% on two stabilizers *K*^+^ and *K*^+^ + PDS respectively in terms of Pearson correlation metric.
- Additionally, we apply our G4-Attention model to predict G4 propensity along cell lines. For a fair comparison with the baseline model named DeepG4, we incorporated chromatin accessibility measures as an additional feature to
- G4-Attention. Both model vanilla G4-Attention and G4-Attention with chromatin accessibility achieve an improvement of 1.37% and 0.55% over DeepG4 (vanilla) and DeepG4 with chromatin accessibility respectively along different cell lines.
- In addition, we apply G4-Attention which is trained to the human-genome dataset on 11 non-human genome datasets and our model achieves benchmark performance over the baseline model G4Detector for the G4 propensity prediction task. Particularly, G4-Attention improves the overall performances of G4Detector by 9.99% and 28.27% on two stabilizers *K*^+^ and *K*^+^ + PDS respectively.
- We also apply our model G4-Attention on held-out datasets from human and other species and our model G4-Attention produces a state-of-the-art model for G4 mismatch score prediction.
- To investigate the robustness of our model, we apply it to highly class-imbalanced scenarios and the experimental results show that G4-Attention produces state-of-the-art results against the second-best model, G4NN.
- Finally, we applied unique visualizations to highlight important features identified by G4-Attention in order to deduce insights underlying the G4-formation mechanism.

## Materials and methods

### Datasets

In this section, we will discuss about the datasets used to train and evaluate our proposed model G4-Attention. Datasets and their problem type used in this work has been depicted in Table 1.

**Table 1.** Overall dataset statistics.

| Dataset Name | Problem Type | Imbalanced |
| --- | --- | --- |
| G4-seq<sub>B</sub> | Classification | x |
| G4-seq<sub>M</sub> | Regression | x |
| G4-seq<sub>Cell</sub> | Classification | x |
| G4-seq<sub>IB</sub> | Classification | ✓ |

#### G4-seq_B_

In the present study, we further extend the application of our model G4-Attention to a balanced dataset, initially introduced in this work [59]. This dataset covers whole-genome G4 measurements across four species [59] (human, mouse, zebrafish, drosophila, C. *elegans*, saccharomyces, leishmania, trypanosoma, plasmodium, arabidopsis, escherichia coli and rhodobacter)^1^. Here, we focus exclusively on the whole-genome data about humans for training and evaluation of our model. As mentioned in this work [59], we obtained two FASTA files for the *K*^+^ and *K*^+^ + PDS for the human genome, respectively and we mentioned these sets as the positive sets in G4 prediction problem. To generate the negative examples for each positive example, we followed the technique used by Barshai *et al*. [59] and we constructed three datasets.

- Random: for every positive sequence, we obtained an equivalent sequence, matching in length, from a randomly selected coordinate within the human genome.
- Dishuffle: for every positive sequence, we randomized its nucleotide arrangement while maintaining the frequencies of dinucleotides [64].
- PQ: predicted G4s in the genome using the regular expression [*G*^3+^*N* ^1−12^]*G*^3+^, where *N* represents any nucleotide.

For all three sets of data, we consider the genome sequences belonging to Chromosome 1 as the test sets and the remaining sequences as the training sets. The overall dataset statistics are depicted in Table 2. The overall instances present for *K*^+^ and *K*^+^ + PDS, for each of three negative sets (random, dishuffle, and PQ), for each species^1^ is present in S1 Table, which is in the supplementary information.

**Table 2.** Dataset Statistics for G4-seq_B_ dataset.

| | $K^+$ | | $K^+ + \text{PDS}$ | |
| --- | --- | --- | --- | --- |
|  | # Training | # Testing | # Training | # Testing |
| Random | 768,256 | 70,736 | 2,427,160 | 231,026 |
| Dishuffle | 794,186 | 74,330 | 2,510,078 | 242,730 |
| PQ | 784,576 | 75,457 | 1,360,065 | 131,206 |

#### G4-seq_M_

In this present study, we further extend the application of our architecture G4-Attention to a regression dataset, as mentioned in this work [65]. This dataset covers whole-genome G4 measurements over 12 species [66] (human, mouse, zebrafish, drosophila, C. *elegans*, saccharomyces, leishmania, trypanosoma, plasmodium, arabidopsis, escherichia coli and rhodobacter)^1^, publicly available via accession code GSE110582. This dataset contains widely studied model organisms and pathogens of clinical relevance. The experiment covers both forward and reverse DNA strands, and assigns to almost every non-overlapping 15 nt-long bin in each of the 12 genome a mismatch score. The G4 mismatch score is defined as the ratio of the number of mismatched base calls observed under a G4-stabilizing condition compared to control conditions over the length of the complete sequence [66]. Its range is 0% to 100%. Here, we consider G4 structures that form under physiological *K*^+^ conditions were identified as well as G4s stabilized by a combination of *K*^+^ and PDS i.e; *K*^+^ + PDS. We consider the genome sequences belonging to Chromosome 1 as the test sets and the remaining sequences as the training sets. We will denote the *K*^+^ for G4-seq_M_ as 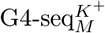 and *K*^+^ + PDS as 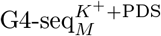 in the upcoming sections for simplicity. The number of sequence scores (mismatch score) in the G4-seq dataset used in our study is summarized in S2 Table, which is in supplementary information.

#### G4-seq_Cell_

In the current work, we also extended the use of our method G4-Attention to G4 ChIP-seq dataset for different cell lines. As Rocher *et al*. [60] mentioned, we downloaded G4 ChIP-seq dataset for HaCaT, K562, and HEKnp cell lines from GEO accession numbers GSE76688, GSE99205, and GSE107690 respectively [54, 67, 68]. We followed the techniques mentioned in Rocher *et al*. [60] to map every cell line to hg19 and to merge peak calling. Again we downloaded G4 ChIP-seq peaks which are already mapped to hg19 for A549, H1975, 293T, and HeLa-S3 cell lines from GEO accession number GSE63874 [69]. We download processed G4-seq peaks mapped to hg19 from GEO accession number GSE63874 [53]. As mentioned in Rocher *et al*. [60], we consider G4-seq from the sodium (Na^+^) and potassium (K^+^) ion conditions and no filtering steps are performed on peak selection.

We adapt the technique used in Rocher *et al*. [60] to create the positive and negative DNA sequences corresponding to each cell line. Positive DNA sequences (active G4 region sequences) are formed by considering G4 ChIP-seq peaks overlapping with G4-seq peaks. We then used the 201-bp DNA sequences centered on the G4 ChIP-seq peaks summits. Negatives sequences (non-active G4 region sequences) are formed by drawing sequences from the human genome with sizes, GC content (%GC) and repeat content (tandem repeat number from Tandem Repeat Finder mask from hg19 genome) similar to those of positive DNA sequences using genNullSeqs function from gkmSVM R package^2^. Additionally, we downloaded processed DNase-seq bigwig files for different cell lines from ENCODE [70] and processed ATAC-seq bigwig files for HaCaT cell lines from GSE7668. Here, we consider the DNA sequences belong to Chromosome 1 for evaluating our model performance and the remaining DNA sequences to train our model.

#### G4-seq_IB_

We acquired human genome named *hg19* /GRCh37 from the the UCSC Genome Browser^3^. We employed the *Python re* package to extract PQS sequences, expanding the conventional definitions of G4. This was achieved using specific regular expression patterns: [*G*^3+^*L*^1−12^]^3+^*G*^3+^, a standard G4 pattern with elongated loop length [71]; [*GN* ^0−1^*GN* ^0−1^*GL*^1−3^]^3+^*GN* ^0−1^*GN* ^0−1^*G*, a enlarged G4 pattern with probable G-run breaks [72]; and [*G*^1−2^*N* ^1−2^]^7+^*G*^1−2^, an uneven G4 pattern [73], where *L* ∈ {*A, T, C, G}* and *N* ∈ *{A, T, C }*. We filtered the redundant and nested G4 sequences by taking into consideration at least one nucleotide. Additionally, we calculated the total number of guanines, ensuring that they are greater than or equal to 12, to facilitate the formation of three-layered G4s. This search was conducted on both strands. For each PQS, the corresponding G4 score was determined by analyzing the experimental peaks and calculating the mean of the continuous signal in the .*bedgraph* format. In the instances where the PQS coordinate lacked an associated signal, the score was set as 0.0. For training purposes, G4 labels were obtained from the G4P ChIP-seq study performed on A549 cell lines, as present in the GEO database with accession number GSE133379. The top 5% of normalized experimental scores were classified as the “positive” class and the remaining are classified as the “negative” class^4^. In this dataset most of the PQS had zero scores, therefore, the classes are highly imbalanced. Thus, we applied our model in a highly imbalanced scenario, which helped us establish the robustness of our model. The distribution of positive and negative classes is shown in Table 3. The dataset was divided into two sub-parts: train and test, where the test subpart includes all the PQS belonging to chromosomes 1, 3, 5, 7, and 9 and the train subpart includes all the PQS belonging to chromosomes 2, 4, 6, 8, 10-22, X, Y. The number of training and testing samples present in the dataset is 1,494,884 and 610,953.

**Table 3.** Distribution of Positive and Negative Samples in the G4-seq_IB_ Dataset.

| Sample Type | Percentage | Number of Samples |
| --- | --- | --- |
| Positive | 5.0% | 105,286 |
| Negative | 95.0% | 2,000,551 |

<sup>2</sup>https://cran.r-project.org/web/packages/gkmSVM.

### G4-Attention Neural Network Architecture

This section starts by outlining the general idea of our proposed neural network framework namely G4 quadruplex prediction by utilizing Attention mechanism (**G4-Attention**) for predicting whether a genome sequence forms G4 or not, followed by a detailed discussion about the different components of our proposed framework. The overview of our proposed architecture is depicted in Fig 2.

**Fig 2.**
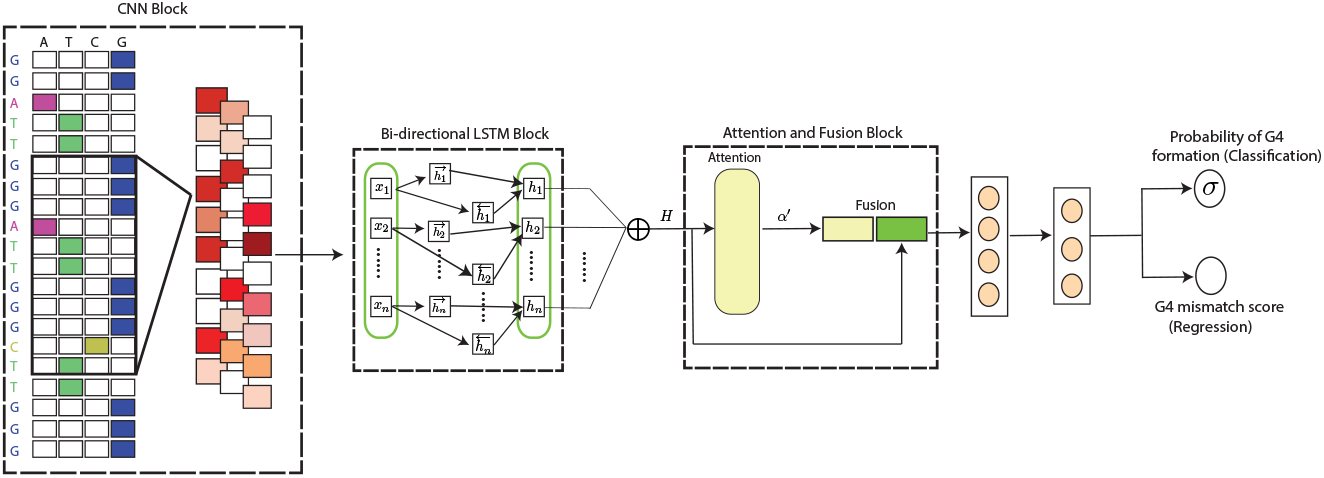
Schematic diagram of our proposed model **G4-Attention**. Our Proposed model has three blocks: CNN block, Bi-LSTM block and finally Attention Fusion block.

#### Encoding of the Genome Sequence

The nucleotide in each position of a genome sequence is represented by a 4-dimensional one-hot vector for training and testing as *A* = (1, 0, 0, 0), *T* = (0, 1, 0, 0), *C* = (0, 0, 1, 0) and *N* = (0, 0, 0, 0). We denote the one-hot vector representation for a sequence as **S**^**O**^ of shape *L* ×4, where *L* is the sequence length. **S**^**O**^ carries the raw information of the sequence.

#### Feature extraction using CNN

We automatically extract local features of **S**^**O**^ using a Conv1D layer. Conv1D layer is a popular architecture in genomics that automatically learns important features from raw sequences [74]. More specifically, the Conv1D layer utilizes **S**^**O**^ as the input raw information of the sequence of shape *n×* 4 and applies a kernel (Θ_*c*_) of shape *K× F* where *K* is the kernel size and *F* is the number of kernels. The output of the CNN is 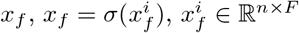, where *σ* is a non-linear function, applied on the intermediate value 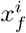 obtained through Conv1D.

#### Feature extraction using Bi-directional LSTM

We pass the learned feature representation of **S**^**O**^ using our Conv1D block, which is *x*_*f*_ to a Bi-LSTM layer, which captures the dependencies between the characters in a sequence of length *n*. We get the output representation 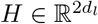, where *d*_*l*_ represents the number of output units used in each LSTM cell. We use Eq. 1 for the execution of the LSTM.

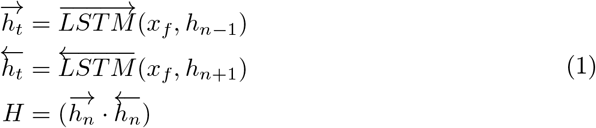

#### Attention and Fusion block of G4-Attention

Here, we use a simplified form of attention mechanism popularly used in natural language processing [75]. The attention block calculates *α* for *H* and the values of *α* quantify the importance of each nucleotide in a genome sequence. We calculate *α* using the Eq. 2.

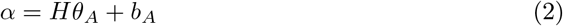

where *θ*_*A*_ is the trainable weight in the attention module, *b*_*A*_ is the bias. *α* is fed into the softmax layer and is normalized to *α*^*′*^ using the following Eq. 3.

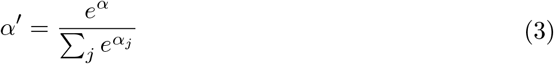

The fusion block receives *H* and *α*^*′*^ from the Bi-LSTM and the attention blocks and subsequently computes the final feature tensor *X* using the Eq. 4.

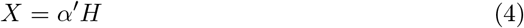

#### Prediction Layer

*X* is fed into two fully connected neural networks and finally, we applied a sigmoid activation function as the task is a binary classification problem. Again, in this study we consider regression task, where we predict a score for a given DNA sequence. Thus *X* is fed into two fully connected neural networks and outputs a single real-valued score, which we denote as a mismatch score. We used the binary cross-entropy loss using the Eq. 5 for the classification task and for the regression task we used the mean squared error (MSE) loss using the Eq. 6 to train the **G4-Attention** model.

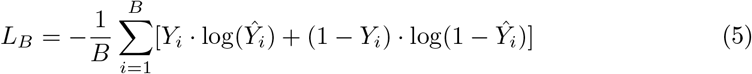

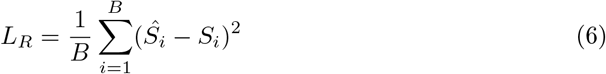

where 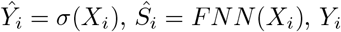 is the class of the *i*^th^ DNA sequence, *S*_*i*_ is the original mismatch score, 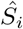 is the predicted mismatch score, *B* is the number of data samples in one batch, *L*_*B*_ is the binary cross entropy loss used for both G4-seq_B_ and G4-seq_Cell_ dataset and *L*_*R*_ is the MSE loss used for G4-seq_M_. Again, in G4-seq_IB_, the number of positive samples is suppressed by the number of negative samples by a large margin, resulting in the network potentially being biased towards the negative samples and failing to correctly identify positive samples when Eq. 5 is considered as the loss function. This issue is tackled by implementing balanced class weights 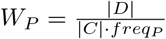 and 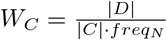 for positive and negative samples respectively. Here |*D*| is the samples |*C*| number of present in the dataset and is the total number of classes. We slightly modify Eq. 5 to introduce class weights in binary classification loss as shown in Eq. 7.

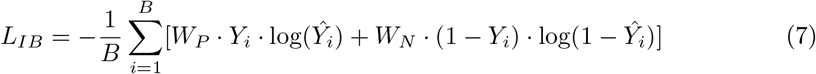

Where *L*_*IB*_ is the weighted binary classification loss used for G4-seq_IB_ dataset, *W*_*P*_ is the class weight for positive class, *W*_*N*_ is the class weight for negative class, *Y*_*i*_ is the original class for a i^th^ DNA sequence and 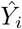 is the predicted class for a i^th^ DNA sequence.

### Parameter Settings

In this work, we fixed the length of the genome sequence to 124 [59] for G4-seq_B_ dataset, 215 [65] for G4-seq_M_ dataset, 201 [60] for G4-seq_Cell_ dataset, and for G4-seq_IB_ we fixed the length of the genome sequence to 1000 [61]. Here, we considered one 1DCNN and one Bi-LSTM layer to train our framework G4-Attention. We trained it for 10 epochs for G4-seq_B_, G4-seq_Cell_, and G4-seq_IB_ datasets and 1 epoch for G4-seq_M_ using Adam [76] optimization with a learning rate of 0.001. We utilized a batch size of 1024 with a seed value of 123 to train our model. We used PyTorch-Ignite^5^ framework for our implementation. One NVIDIA A6000 48GB GPU and one NVIDIA A100 80GB GPU were used to train and evaluate our model G4-Attention.

## Results and Discussion

### Evaluation Criteria

We determine the performance of our proposed model using four widely used metrics; the area under the ROC curve (AUC) for G4-seq_B_ and G4-seq_Cell_, the area under P-R curve (AUPRC) for G4-seq_IB_, Pearson correlation, precision, recall and F-score for G4-seq_M_ using the Eqns. 8, 9 and 10 respectively.

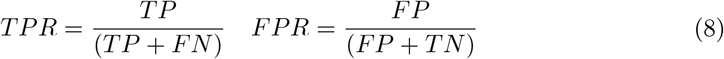

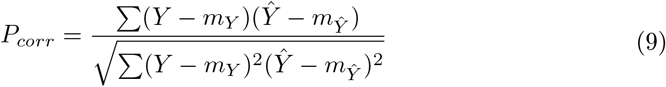

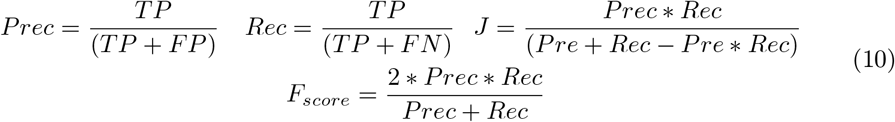

where 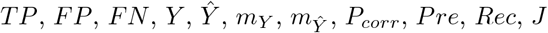 and *F*_*score*_ are true-positive, false-positive, false-negative, original mismatch score, predicted mismatch score, mean of original mismatch score, mean of predicted mismatch score, Pearson correlation, precision, recall, jaccard score and F-score respectively. The AUC measures the performance of the model by calculating the relationship between the false-positive rate (*FPR*) and the true-positive rate (*TPR*) at different probabilities or thresholds. However, AUC is not optimal for imbalanced classification problems [61]; therefore we additionally evaluate AUPRC to determine precision and recall of positive G4 samples in class imbalanced scenarios.

### Research Questions

In this section, we pose a few research questions which are central to our research work.

#### RQ1: Effectiveness of G4-Attention for predicting G4 propensities and mismatch score

How well G4-Attention accurately predict the G4 propensities of a given genome sequence in the balanced scenario? Furthermore, how well G4-Attention predicts the G4 mismatch score and also the G4 propensities from that mismatch score? Also how effectively G4-Attention predicts G4 propensities of a given genome sequence with respect to cell lines?

#### RQ2: Performance of G4-Attention on non-human species

How does our model perform in cross-domain testing, specifically in predicting G4 formation in non-human species? Furthermore, how well G4-Attention predicts G4 mismatch score prediction in non-human species?

#### RQ3: Performance of G4-Attention on class imbalance scenario

What is the effectiveness of our model in scenarios characterized by class imbalance, specifically in situations where the quantity of positive samples is substantially lower than that of the negative samples?

#### RQ4. Interpretability of G4-Attention

How does our model learn key sequence features in G4 formation in predicting G4 propensities and also predicting the G4 mismatch score for a genome sequence?

### Performance of G4-Attention on G4 Propensity Prediction

In relation to the first part of RQ-1, we compare our model with the existing computational methods that are used to predict G4 sequences in the balanced dataset, G4-seq_B_. Here, we make a comparison between our model and 6 state-of-the-art methods, including Quadron [57], pqsfinder [48, 49], G4Hunter [50–52], G4Detector [59], G4Catchall [77] and G4Boost [78]. All methods are applied in identical experimental conditions and utilize the optimal parameters as specified in their respective research articles.

#### G4Hunter

This method utilizes domain knowledge-based rules, using a sliding window approach to determine G4 propensity by analyzing the GC content in a specific sequence.

#### pqsfinder

This method identifies G4s by locating G-tracts in the genome sequence. **Quadron:** In this approach, the objective is to predict the robustness of the DNA G4 structure utilizing machine learning.

#### G4Detector

In this approach, the CNN-based architecture incorporates both raw sequence data and RNA secondary structure information as input.

#### G4Catchall

In this algorithm, a regular expression-based technique is used to predict G4 propensities given in input DNA sequences.

#### G4Boost

In this method, a decision tree-based machine learning algorithm is used to predict the G4 propensities of a given DNA sequence.

Fig. 3 (a) and (b) we report the AUC score (the higher the AUC, the better the model) using Eq. 8 for G4 propensity prediction task on three negatives types on two stabilizers *K*^+^ and *K*^+^ + PDS. We observe that our proposed architecture G4-Attention produces current state-of-the-art (SOTA) results on three negative types on two stabilizers *K*^+^ and *K*^+^ + PDS. The network trained solely on sequence information exhibited notable performance on the *K*^+^ dataset, achieving AUC scores of 0.989, 0.981, and 0.999 for random, dishuffle, and PQ negatives respectively.

**Fig 3.**
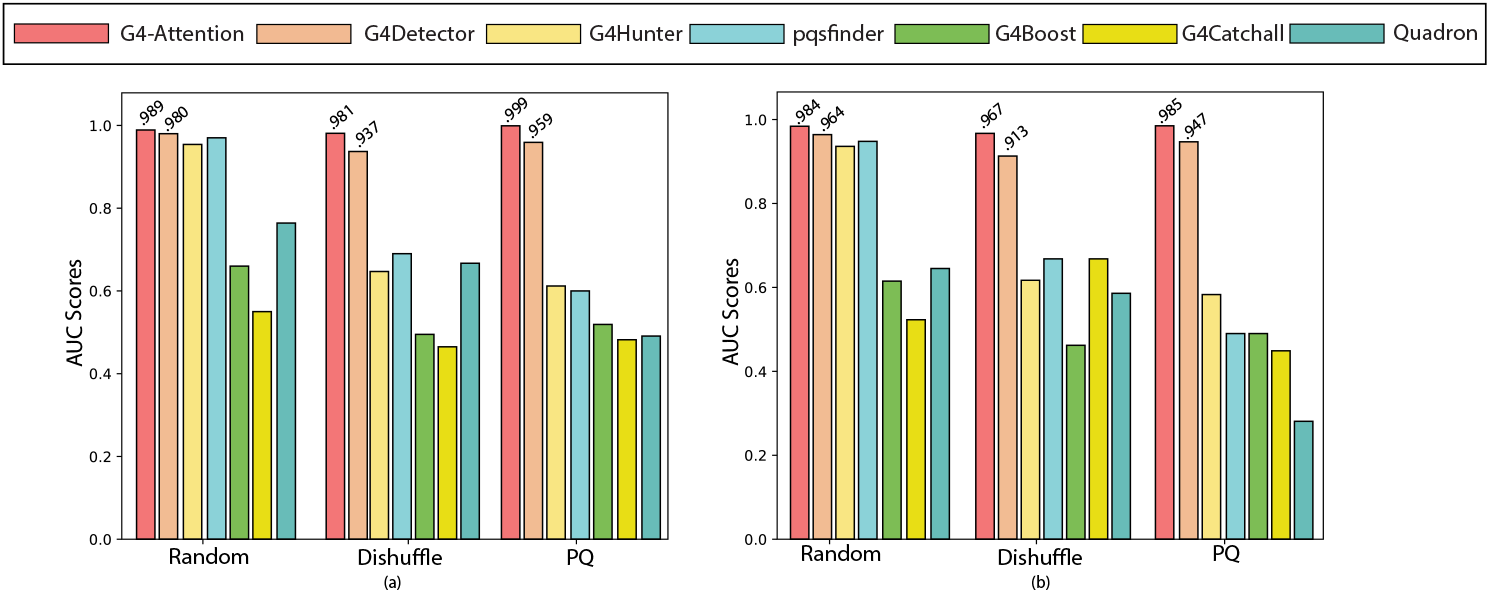
G4-Attention outperforms all the existing techniques across all negative types (a) *K*^+^ and (b) *K*^+^ + PDS on held out chromosome 1 on G4-seq_B_ dataset. Only, G4-Attention result and the second best model result is shown at the top of the bars.

Specifically, the percentage of improvement against the second best model which is G4Detector in random, dishuffle, and PQ negatives on *K*^+^ stabilizer are 0.92%, 4.70%, and 4.17% respectively. The average improvement is 3.26%. We also compare our model G4-Attention performance against other baselines G4Hunter, pqsfinder, Quadron, G4Catchall, and G4Boost. We observe from Fig. 3 (a) that our model G4-Attention produces an improvement of 4.17% against G4Detector in the PQ negative set, which indicates that G4-Attention performs better than G4Detector in the most challenging of the three proposed classification problems [59]. The negative set of PQ is formulated from genomic regions characterized by a G-rich regular expression, thereby demonstrating the capability of our method to predict G4 formation in genomic segments populated with G-rich sequences [59]. Furthermore, we observe from Fig. 3 (b) that G4-Attention produces remarkable AUC scores of 0.984, 0.967, and 0.985 for random, dishuffle, and PQ negatives on *K*^+^ + PDS stabilizer respectively, which is the current SOTA results. Specifically, the percentage of improvement against G4Detector (second best model) is 2.07%, 5.91%, and 4.01% for random, dishuffle, and PQ negatives on *K*^+^ + PDS stabilizer respectively. The overall average improvement is 3.99%.

### Performance of G4-Attention on G4 Mismatch Prediction

To address the second part of RQ-1, we compare G4-Attention’s performance with the existing algorithms that are used to predict the G4 mismatch score and identify G4-forming sequence i.e; G4 propensities by utilizing G4 mismatch score. Here, we tested the ability of G4-Attention to predict G4 mismatch score against trained models of 10 SOTA algorithms including G4mismatch [65], G4Hunter [50–52], pqsfinder [48, 49], g4predict [79], Quadron [57], PENGUINN [58], DeepG4 [60], G4Catchall [77] and G4Boost [78]. A brief description for G4Hunter, pqsfinder, Quadron, G4Catchall, and G4Boost are present in the section “Performance of G4-Attention on G4 propensity prediction” and for the other models we will discuss in this section.

#### G4mismatch

This method utilizes 1D CNN-based architecture which incorporates raw DNA sequence as input and is used to predict the G4 mismatch score. **g4predict:** This method utilizes a regular expression-based algorithm that takes raw DNA sequence as input to generate a score as mentioned in Parker *et al*. [79].

#### PENGUINN

This algorithm uses a 1D CNN-based method which takes raw DNA sequence via randomized mutation technique and generates a score between 0 and 1.

#### DeepG4

This algorithm also uses a 1D CNN-based method along with chromatin accessibility of DNA as an extra feature to predict G4 propensities of a raw DNA sequence.

We take G4Hunter with a window size of 25 and a threshold of 1, as mentioned in Bedrat *et al*. [50]. We ran pqsfinder and g4predict with their parameters as mentioned in the works [48, 79] by setting a threshold to 45 for pqsfinder, as it was found to be the optimal threshold for identifying G4 propensities [48, 49]. For Quadron, we set a threshold of 25 to match the 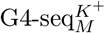 dataset and 35 for 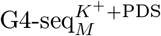 dataset (we did not use the threshold 19 as mentioned in the paper [57] as it produces poor results). For PENGUINN, we used a genome-wide scanner to predict a score for every 200 nt-long sequence in chromosome 1, similar to the scan performed in the work G4mismatch [65]. We follow Barshai *et al*. [65] to use two different thresholds (as defined in the original work [58]) 0.5 and 0.85 denoted by PENGUINN(s) and PENGUINN(q), respectively, to filter the scan result. For DeepG4, we used the parameters mentioned in the study [60]. Similarly, for G4Catchall and G4Boost, we follow the original studies [77, 78] to set the parameters. Finally, for G4mismatch, we follow the optimal parameter mentioned in the study [65]. All methods were fed human genomes belongs to chromosome 1 from G4-seq_M_ dataset to predict mismatch score and then identify G4-forming sequence (G4 propensities) and uses human genomes belongs to other chromosomes as to train all the algorithms mentioned above.

### G4-Attention prediction performance on mismatch score prediction

We tested G4-Attention on the held-out test, which is chromosome 1 of human both for 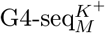 and 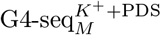. Since both the datasets 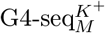 and 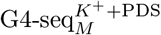 are regression datasets, we take Pearson Correlation score as evaluation metric as shown in Eq. 9 to measure the performance of our model G4-Attention against the baselines. Note that, a higher Pearson Correlation score means more improvement of the model’s performance. Also for the baseline models, Pearson Correlation is used as evaluation metrics [65]. In Figs. 4 and 5, we report Pearson Correlation scores for 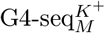 and 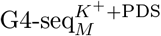 respectively. We observed that G4-Attention achieves a Pearson Correlation score of 0.87 for 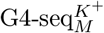 and the second best performing model, which is G4mismatch produces a Pearson Correlation score of 0.84. The average improvement for *K*^+^ against the second-best model, G4mismatch is 3.57% which is quite impressive. For 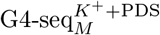, G4-Attention also produces a Pearson Correlation score of 0.94, but the second best model which is G4mismatch produces a mismatch score of 0.91. The average improvement for 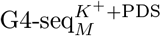 against the second-best model is 3.30% which is also a significant improvement over the second-best performing model.

**Fig 4.**
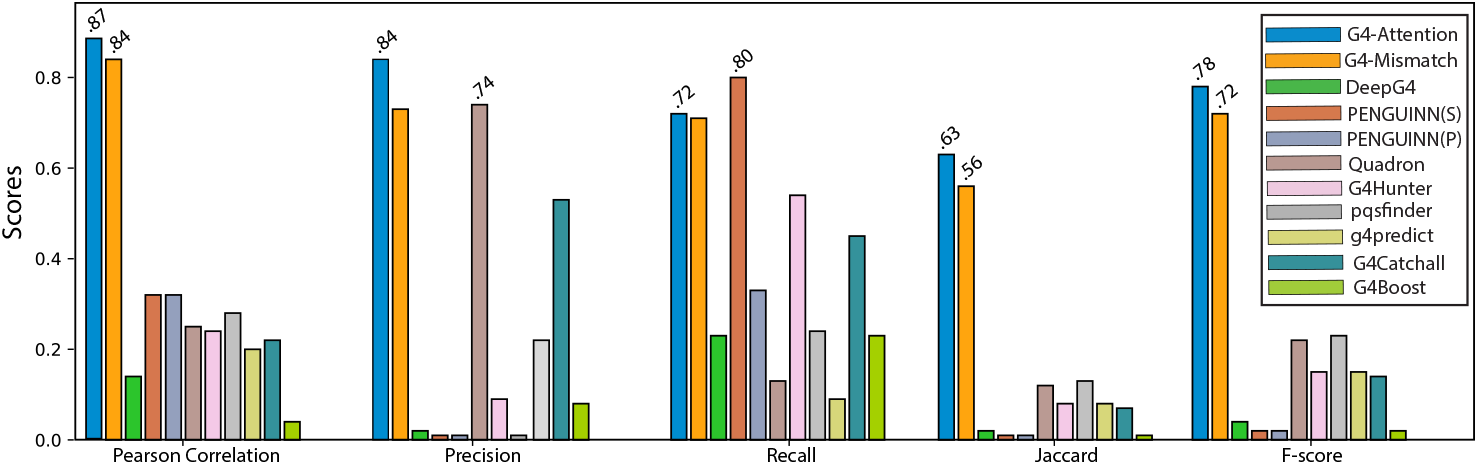
G4-Attention outperforms all the existing baselines across *K*^+^ stabilizer on held out chromosome 1 on G4-seq_M_ dataset. Only, G4-Attention result and the second best model result is shown at the top of the bars.

**Fig 5.**
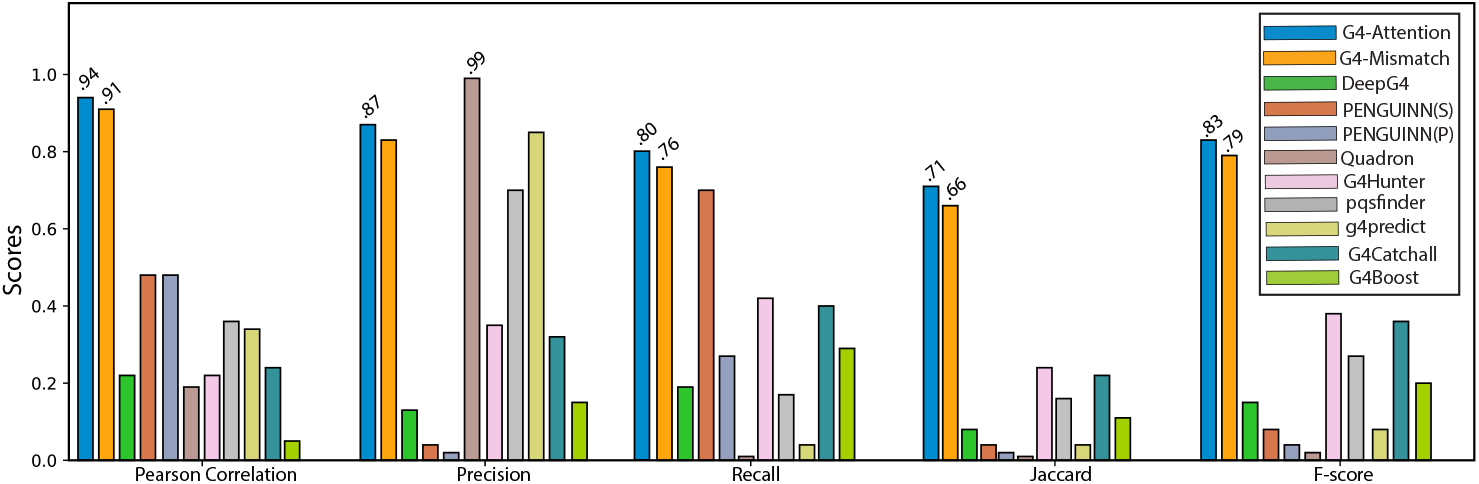
G4-Attention outperforms all the existing baselines across *K*^+^ + PDS stabilizer on held out chromosome 1 on G4-seq_M_ dataset. Only, G4-Attention result and the second best model result is shown at the top of the bars.

### Comparison of G4 Propensity Prediction Performance

We next utilize G4-Attention predictions to identify G4 propensities from the predicted mismatch score. To test the G4 detection ability, we used G4-Attention to predict mismatch score to all genomes belonging to chromosome 1 i.e.; the held-out test set using dataset G4-seq_M_ for both stabilizers 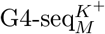 and 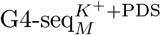. From the predicted scores of chromosome 1, we identified G4 hit regions. We used the threshold mentioned in the work G4mismatch [65], which is 25 for 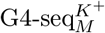 and 35 for 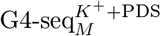 to assign binary label to each one of the genome sequences, 1 for a genome sequence identified as G4 and 0 otherwise. G4s identified by the G4-seq method [66] in chromosome 1 were used as ground truth and labeled similarly. Since this is a classification task, we used precision, recall, Jaccard, and F-score as evaluation metrics which is shown in the Eqs. 10. Note that increasing of precision, recall, Jaccard, and F-score indicates more improvement in the model’s performance.

In Fig. 4, we report the precision, recall, Jaccard, and F-score for 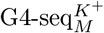. We see that Quadron presents slightly higher precision than G4mismatch (second best model for Pearson Correlation score), but G4-Attention produces a precision score of 0.84 which is 13.51% improved than Quadron. On the other hand, for recall, PENGUINN(s) shows higher recall than our model G4-Attention, scoring 0.80 on 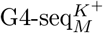 dataset but scoring a low precision score of 0.01. For the Jaccard score, G4-Attention produces the best result of 0.63 which is a 12.50% improvement over the second best method G4mismatch for 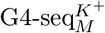 dataset. Finally, for the F-score, G4-Attention produces a SOTA result which is 0.78. More specifically this result is a 8.33% improvement over the second best performing method G4mismatch on 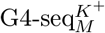 dataset.

In Fig. 5, we report the precision, recall, Jaccard, and F-score for 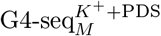. For precision, Quadron produces a precision score of 0.99, which outperforms G4-Attention precision score which is 0.87. An obvious reason could be that Quadron is originally trained on *K*^+^ + PDS stabilized PQs, but Quadron produces a poor recall score of 0.01. For recall, our model G4-Attention produces the best score among all other baseline models. More specifically, it produces a recall score of 0.86 which is a 5.26% improvement over the second-best model G4mismatch. For Jaccard and F-score, G4-Attention performs best among all the baseline models. Specifically for Jaccard, G4mismatch produces a score of 0.71 which is a 7.58% improvement than the second best performing method G4mismatch and for F-score, G4mismatch produces a score of 0.83 which is a 5.06% improvement than the second best model G4mismatch.

### Performance of G4-Attention on G4 Identification Along Cell Lines

In this section, we discuss our third part of the RQ-1 to understand whether our proposed method G4-Attention performs accurately along the cell lines. Here, we slightly modified our proposed method G4-Attention to incorporate chromatin accessibility values as additional features, we denote this model as G4-Attention(WA) and the model without chromatin accessibility i.e.; the vanilla version of G4-Attention as G4-Attention(WoA). DNA sequence belonging to chromosome 1 has been considered as a test set for the HaCaT cell line and the sequence belongs to the remaining chromosome to train both G4-Attention(WoA) and G4-Attention(WA). We take chromatin accessibility values considering the G4-seq_Cell_ dataset. In Fig 6 (a), we report the performance of G4-Attention without using accessibility as additional features (G4-Attention(WoA)) and (b) we report the performance of G4-Attention with accessibility as additional features (G4-Attention(WA)) along the cell lines used for prediction purpose. Here, we take AUC scores, present in Eq. 8 to measure the performance of G4-Attention.

**Fig 6.**
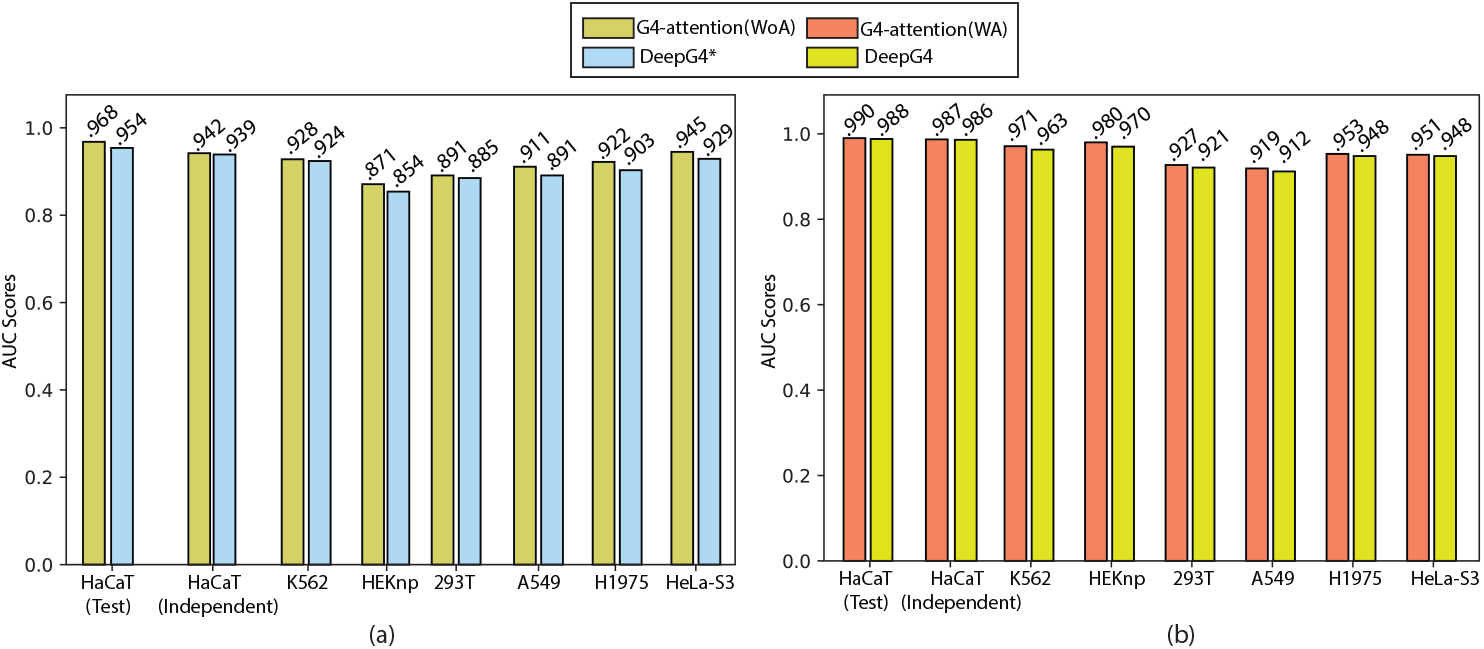
Performance of predictions of active G4 regions by G4-Attention(WoA) and G4-Attention(WA) along the cell lines which is present in x-axis. Here DeepG4* is the DeepG4 with out using chromatin accessibility as additional features. AUC scores for G4-Attention(WoA), G4-Attention(WA), DeepG4* and DeepG4 are indicated at the top of the bars.

From Fig 6 (a), we see that G4-Attention(WoA) obtained excellent predictions of active G4 regions from HaCaT cells on the testing set producing an AUC score of 0.968 which outperforms the DeepG4* result by 1.47%. On an independent ChIP-seq experiment done with the same cell line (GEO GSE99205 accession), G4-Attention(WoA) outperforms DeepG4* by a margin of 0.32%. Overall, G4-Attention which was trained using HaCaT cell line data by considering the dataset G4-seq_Cell_, could well predict in other cell lines. For instance, we see that, for K562, G4-Attention(WoA) achieves a high AUC score of 0.928 which outperforms the DeepG4* result by 0.43%. For HEKnp, 293T, A549, H1975, and HeLa-S3 we see that G4-Attention(WoA) produces an impressive AUC score of 0.871, 0.891, 0.911, 0.922, and 0.945 respectively. These results outperforms the DeepG4* result by 1.99%, 0.68%, 2.24%, 2.10% and 1.72%.

As mentioned in the work, [60] chromatin accessibility could help to produce cell-type specific predictions. From Fig 6 (b), we observe that adding chromatin accessibility significantly increases cell-type specific prediction accuracy. For instance, the AUC score of HaCaT (test) and HaCaT (independent) were 0.990 and 0.987 for G4-Attention(WA) as compared to 0.968 and 0.942 for G4-Attention(WoA). Also G4-Attention(WA) outperforms DeepG4 results by 0.20% and 0.10% on HaCaT (test) and HaCaT (independent) cell lines. Again, for K562, HEKnp, 293T, A549, H1975, and HeLa-S3, we observe that G4-Attention(WA) (K562: AUC = 0.971; HEKnp: AUC = 0.980; 293T: AUC = 0.927; A549: AUC = 0.919; H1975: AUC = 0.953; HeLa-S3: AUC = 0.951) produces better AUC than G4-Attention(WoA) (K562: AUC = 0.928; HEKnp: AUC = 0.871; 293T: AUC = 0.891; A549: AUC = 0.911; H1975: AUC = 0.922; HeLa-S3: AUC = 0.945). Also for all of these cell lines, G4-Attention(WA) outperforms DeepG4 by a percentage of improvement for K562 (0.83%), HEKnp (1.03%), 293T (0.65%), A549 (0.77%), H1975 (0.53%) and HeLa-S3 (0.32%).

Thus, all these results collectively demonstrated the ability of G4-Attention(WoA) and G4-Attention(WA) to predict cell-type specific G4 propensities accurately from DNA sequences and chromatin accessibility. Moreover, results also revealed the importance of incorporating chromatin accessibility into G4-Attention for cell-type specific predictions.

### Performance of G4-Attention in Non-Human Species on G4 propensity prediction

To address the first part of the RQ-2, we applied our model G4-Attention on the multiple species dataset reported by Barshai *et al*. [59], which provides a perfect test set for G4-Attention. This test set contains DNA sequences only for three species other than humans (*H. sapiens*). These are mouse (*M. musculus*), zebrafish (*D. reiro*), and drosophila (*D. melanogaster*) which is publicly available via accession code GSE110582. But, in GSE110582, there are other 7 species which is not used in this work [59]. Thus, we follow the techniques mentioned in Barshai *et al*. [59] to create the test dataset for the other 7 species. These species are *C. elegans, Saccharomyces cerevisiae* (*S. cerevisiae*), *Leishmania major* (*L. major*), *Trypanosoma brucei* (*T. brucei*), *Plasmodium falciparum* (*P. falciparum*), *Arabidopsis thaliana* (*A. thaliana*), *Escherichia Coli* (*E. Coli*) and *Rhodobacter sphaeroides* (*R sphaeroides*). For these 7 species, we create three negative sets (random, dishuffle, and PQ) as mentioned in G4Detector [59]. Here, we measured the ability of our model G4-Attention in a cross-domain testing scenario. We utilized G4-Attention, trained on the human genome, to conduct zero-shot testing on a range of non-human species, none of which were included in the initial training dataset. In Fig. 7, we report the zero-shot prediction performance of G4-Attention on 11 species in terms of AUC score, which is present in the Eq. 8.

**Fig 7.**
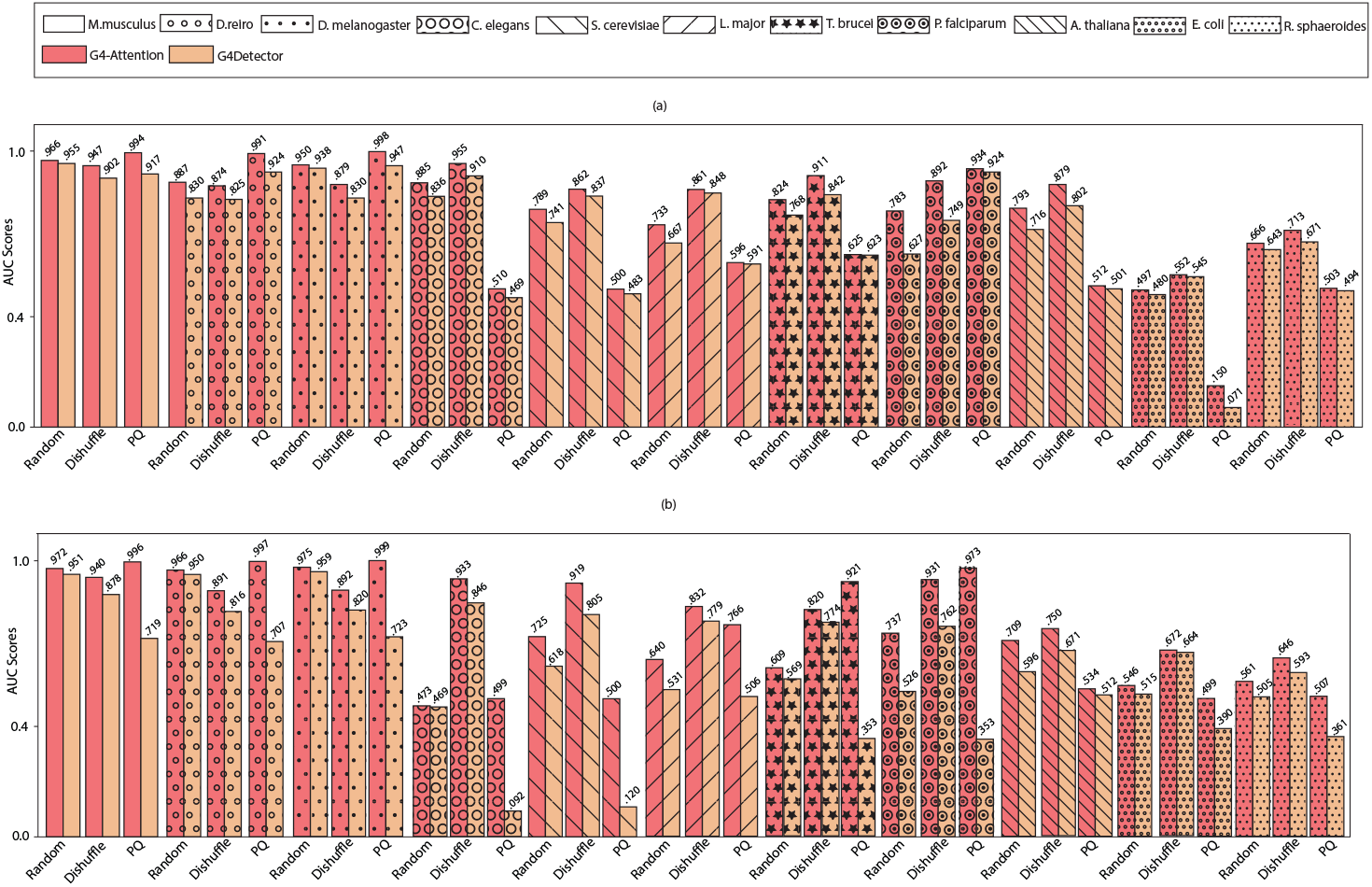
The comparative analysis of G4-Attention’s performance on novel datasets from eleven distinct species against G4Detector is illustrated in the figure. This evaluation specifically focuses on the AUC scores on (a) *K*^+^ datasets and (b) *K*^+^ + PDS datasets present in G4-seq_B_. AUC scores for G4-Attention and G4Detector are indicated at the top of the bars.

From Fig. 7, we observed that G4-Attention produces current better results in the cross-domain test data. More specifically, Fig. 7 (a) shows that G4-Attention produces excellent prediction performance for non-human species like mouse, zebrafish, drosophila, etc. on stabilizer *K*^+^ present in G4-seq_B_ dataset. Here we take G4Detector [59] as the baseline model to compare the performance of G4-Attention in the cross-domain testing as it achieves current SOTA result on the cross-domain testing. In particular, for random negative set, G4-Attention achieves AUC score of 0.966, 0.887, 0.95, 0.885, 0.789, 0.733, 0.824, 0.783, 0.793, 0.497 and 0.66 respectively, which improves the AUC scores of G4Detector by 1.15%, 6.87%, 1.28%, 5.86%, 6.48%, 9.89%, 7.29%, 24.88%, 10.75%, 3.54% and 3.62% respectively for 11 species. The average improvement obtained by G4-Attention over G4Detector on random negative set is 9.99%. As mentioned in Barshai *et al*. [59], classification on random genomic set is simplest among all three negative sets, the average improvement is remarkable as already G4Detector predicts good AUC score and top of that G4-Attention predict better AUC as we see from the Fig. 7 (a). For dishuffle negative set, G4-Attention produces AUC score of 0.947, 0.874, 0.879, 0.955, 0.862, 0.861, 0.911, 0.892, 0.879, 0.552, and 0.713 respectively, which improves the AUC score of G4Detector by 4.99%, 5.94%, 5.90%, 4.95%, 2.99%, 1.52%, 8.19%, 19.09%, 9.60%, 1.28% and 6.26% respectively for 11 species. The average improvement obtained by G4-Attention over G4Detector on dishuffle negative set is 6.42%. In dishuffle negative set, the sequence preserves the same nucleotide sequence but lacks the correct ordering form a G4 [59]. This is the main reason that the average improvement is lesser than random negative sets. In PQ negative set, G4-Attention produces AUC score of 0.994, 0.991, 0.998, 0.501, 0.500, 0.596, 0.625, 0.936, 0.512, 0.150 and 0.5034 respectively, which improves the AUC score of G4Detector by 8.39%, 7.25%, 5.39%, 6.82%, 3.52%, 3.52%, 0.85%, 0.32%, 1.33%, 2.20%, 111.27%, 1.90% respectively. The average improvement obtained by G4-Attention over G4Detector on the PQ negative set is 13.57%, which is a remarkable improvement, as classifying genome sequence as G4 in PQ negative is quite challenging. The main reason is that PQ negative set is constructed out of areas in the genome containing a G-rich regular expression (subsection “Datasets” in section “Materials and methods”), thus we observe that there are some similarities between the PQ negative set and the G4 positive set, which most probably contains similar G-rich expressions [59].

Again, Fig .7 (b) shows that G4-Attention produces SOTA prediction performance for non-human species on stabilizer *K*^+^ + PDS present in G4-seq_B_ dataset. We choose G4Detector [59] as the baseline model to compare G4-Attention performance. Similar to *K*^+^, we have three negative sets, random, dishuffle, and PQ for *K*^+^ + PDS. For random, G4-Attention produces AUC scores of 0.972, 0.966, 0.975, 0.473, 0.725, 0.640, 0.609, 0.737, 0.709, 0.546, and 0.561 respectively, which improves the AUC score of G4Detector by 2.21%, 1.68%, 1.67%, 0.94%, 17.28%, 20.61%, 7.02%, 40.11%, 18.96%, 18.95%, 6.00%, and 11.56%. The average improvement is 11.59%, which is similar to a random set of *K*^+^ as predicting in random negative set is easier, and thus G4Detector also produces a good AUC score. For dishuffle, G4-Attention produces AUC score of 0.940, 0.891, 0.892, 0.933, 0.919, 0.832, 0.820, 0.931, 0.750, 0.672 and 0.646 respectively, which improves the AUC score of G4Detector by 7.06%, 9.19%, 8.78%, 10.28%, 14.21%, 6.80%, 5.82%, 22.18%, 11.79%, 1.19% and 8.94% respectively. The average improvement is 7.97%. From *K*^+^, we know that prediction performance is decreasing in dishuffle negative set, similar kind of things happens for *K*^+^ + PDS. Finally, for PQ negative set, G4-Attention produces AUC score of 0.996, 0.997, 0.999, 0.499, 0.500, 0.766, 0.921, 0.973, 0.534, 0.499 and 0.507 respectively, which improves the AUC score of G4Detector by 38.52%, 41.02%, 38.17%, 441.80%, 316.67%, 51.27%, 161.22%, 176.03%, 4.34%, 28.61% and 40.36% respectively. The average improvement is 65.25% over G4Detector prediction, which is notable as classification of G4 genome sequence in PQ negative set is challenging [59].

### G4-Attention Prediction Performance Evaluation on Inter-species

In this section, we will discuss the second part of our RQ-2. Here, we follow the work by Barshai *et al*. [65] to apply G4-Attention in Non-Human Species on G4 mismatch score prediction. Similar to Barshai *et al*., [65] we took the recently published G4-seq datasets of 12 species to test if the G4-folding principle learned from one species can transform to other species. We trained G4-Attention on each of the species and tested each species-specific model on all other species-specific datasets. We report the species-specific performance of G4-Attention using 12 species in the Fig. 8.

**Fig 8.**
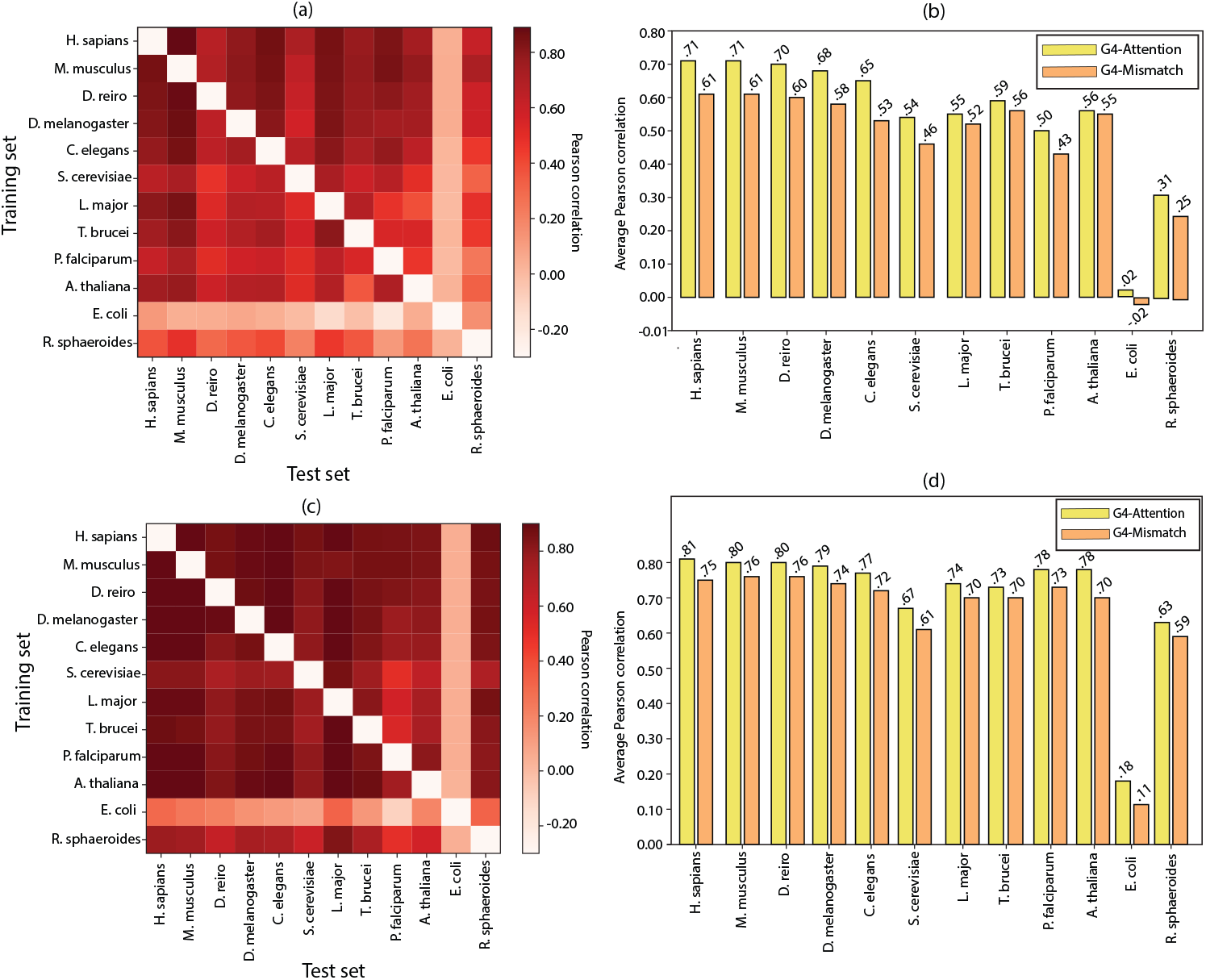
Inter-species G4-Attention (a) *K*^+^ and (c) *K*^+^ + PDS prediction performance. Each model was trained on data of one species obtained from (a) 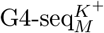 and (c) 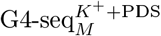 and tested on all other species both for (a) 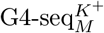 and (c) 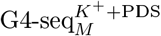. Prediction performance is reported in Pearson correlation of predicted and measured G4 mismatch score. The average Pearson correlation between G4-Attention and G4mismatch for (b) 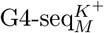 and (d) 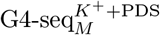 on data of each species.

Fig. 8 (a) shows mismatch score prediction performance in terms of Pearson correlation on stabilizer *K*^+^ using 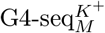 dataset and Fig. 8 (b) shows the comparison of average Pearson correlation for each species between our model G4-Attention and G4mismatch [65]. From Fig. 8 (b), we see that human, mouse, and zebrafish models (G4-Attention) exhibit the best average performance, achieving an average Pearson correlation of 0.71, 0.71, and 0.70 which outperforms human, mouse and zebrafish models (G4mismatch) by 16.49%, 16.51%, and 17.78%, on the other 11 species for each model trained on the 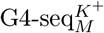. Surprisingly, the Leishmania-specific model achieved excellent performance when predicting mismatch scores in other species, even in some cases outperforming species, which are phylogenetically more related [65]. For example, Leishmania-specific model (G4-Attention) achieved a Pearson correlation of 0.80, which outperforms Leishmania-specific model (G4mismatch) achieved a Pearson correlation of 0.72 on the human 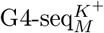 dataset, and also Leishmania-specific model (G4-Attention) outperforms *C. elegans* model (G4mismatch) as it achieves a Pearson correlation score of 0.66 on the human 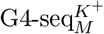 dataset. In contrast, for some species, G4-Attention shows below-par performance but produces better performance than G4mismatch models. For example, the bacteria Rhodobacter model (G4-Attention) achieved an average Pearson correlation of 0.31 which outperforms the mean Pearson correlation of the Rhodobacter model (G4mismatch) by 22.55%. We observe that *E. coli* -specific model (G4-Attention) performs worst, as it produces a very low mean Pearson correlation of around 0.02, which is better than the Pearson correlation of *E. coli* -specific model (G4mismatch) which is -0.02. A recent study revealed a unique and non-random localization of G4-forming sequences in bacterial genomes, which may partially explain the performance of the bacterial-specific models [80].

Fig. 8 (c) shows mismatch score prediction performance in terms of Pearson correlation on stabilizer *K*^+^ + PDS dataset and Fig. 8 (d) shows the comparison of average Pearson correlation for each species between our G4-Attention and G4mismatch. A similar type of result has been obtained for the *K*^+^ + PDS dataset, as we previously got for the *K*^+^ dataset.

### Model Comparison in Imbalanced Scenario

In this section, we turned our attention to RQ-3 to understand whether our proposed method G4-Attention performs accurately in practical situations i.e., for the negatively skewed dataset. The occurrence of G4 structures within the human genome exhibits a pronounced negative skewness. The G4-seq experiment revealed approximately 1,400,000 G4s distributed throughout the human genome [66]. Given the human genome’s composition of over six billion nucleotides, the frequency of G4 can be estimated to occur approximately every 4,200 to 15,000 nucleotides. This drives the necessity to assess the efficacy of G4-Attention when applied to a dataset with a negative skewness. Due to the class imbalance, we measured the performance of G4-Attention using the AUPRC, in addition to AUC.

We use the class imbalanced dataset named G4-seq_IB_ and measure the performance of G4-Attention against the baseline model G4NN^6^ [61], a ResNet [62] based architecture. From Table 3 we observe that the dataset G4-seq_IB_ consists of high negative skewness. To address this problem we used Eq. 7 as the loss function to optimize the model parameters of the G4-Attention. From Fig. 9 we observe that it produces AUC and AUPRC scores of 0.986 and 0.771 respectively, which outperforms the existing model G4NN [61] by an improvement of 1.44% and 31.57% respectively. This finding establishes that our model works equally well in negatively skewed datasets, thereby proving the robustness of our model.

**Fig 9.**
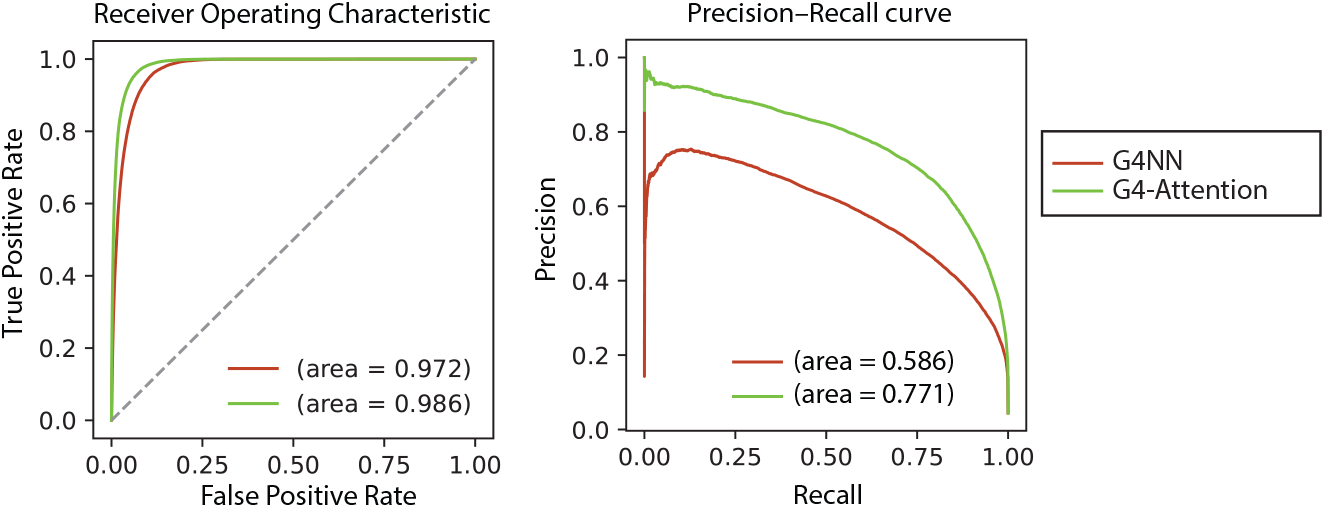
The performance of G4-Attention on test chromosomes 1, 3, 5, 7, 9 in the G4-seq_IB_ dataset is depicted in this figure, where both AUROC and AUPRC are present.

### Interpretation of G4-Attention for Learning Key Sequence Features in G4 Propensity Prediction

Here, we address the first part of RQ-4. To address RQ-4, we follow the Barshai *et al*. [59] and we tried to determine how our model G4-Attention learns important features in G4 propensity prediction. The key features identified by G4-Attention in a specific sequence were visualized using the integrated gradient (IG) method and a mutation map approach. In this study, we take the genome sequence used in the work G4Detector [59]. We calculated attribution score and mutation sensitivity scores using the three *K*^+^ (interpretation results for three negatives for *K*^+^ + PDS stabilizers present in the supplementary information) models, by considering a unique negative set (random, dishuffle and PQ) is shown in Fig. 10.

**Fig 10.**
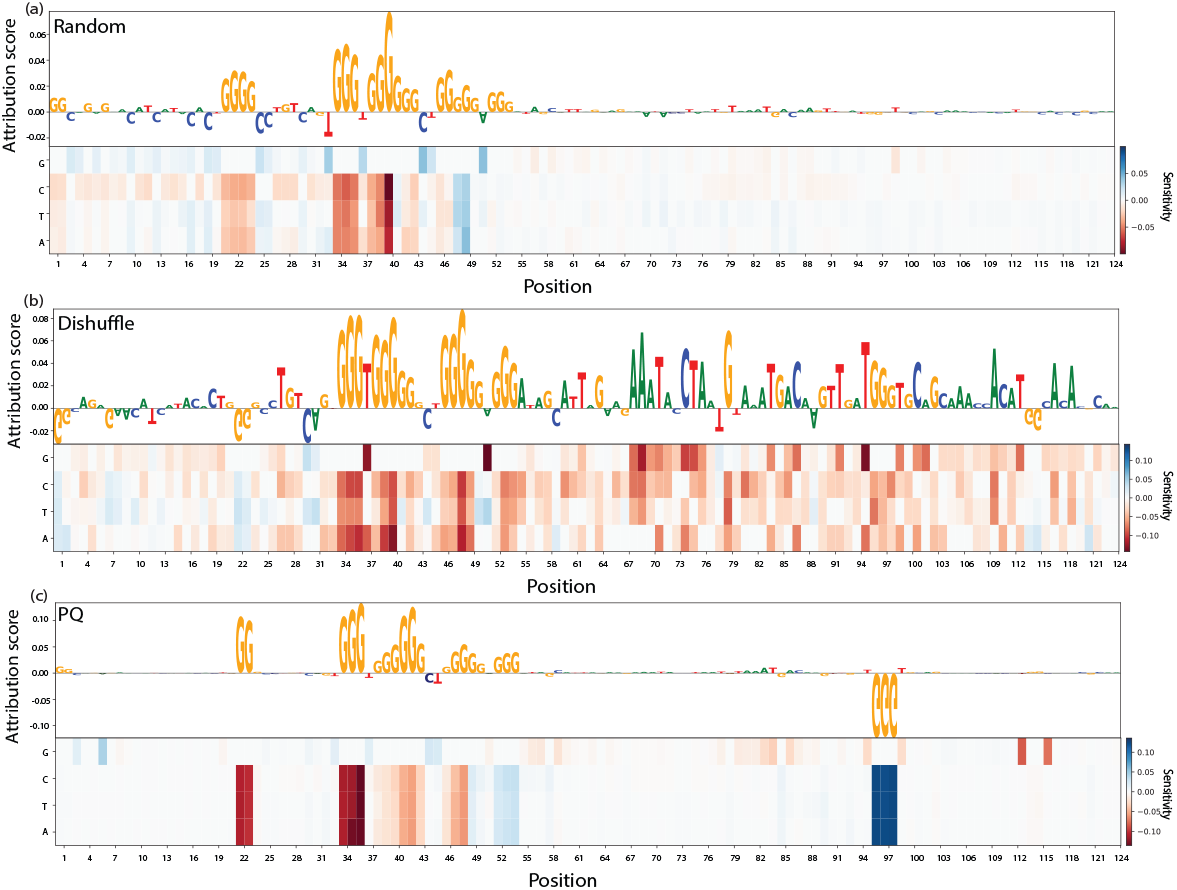
An illustration of attribution scores determined through the integrated gradient method and mutation maps for a specific sequence, using models that have been trained with the G4-seq_B_ dataset for a single genome sequence. This example is depicted in figures (a), (b), and (c), demonstrating that stretches of G’s, which are separated by short loops, tend to get relatively high scores in models trained on the Random, Dishuffle, and PQ negative collections.

The attribution scores indicate that the models trained on the random, dishuffle negative sets are based on G-rich sub-sequences. The relevant G4s reside in continuous stretches. For the models trained on the random and dishuffle negative sets, higher attribution scores were given to pairs of guanine stretches separated by short connecting strands, which are shown to exhibit greater stability [71] which is depicted in Fig.10 (a) and (b). More specifically, we observe that G4-Attention assigned greater importance to inner Gs compared to outer Gs. The importance of inner Gs compared to outer Gs was recently reported in an independent analysis of the conservation of human G4 sequences [81]. We observe that our result is similar kind of the results observed for random and dishuffle in the work G4Detector [59] which also signifies the results present in Fig. 3 (a) results observed for random and dishuffle in the section “Performance of G4-Attention on G4 Propensity Prediction”. But for PQ, negative set G4Detector performs worst, as we see from G4Detector that G-rich genomic areas received negative attributions which are also present in Fig. 3 (a). In that contrast, G4-Attention trained on PQ negative set, G-rich genomic areas received positive attributions, which gives an interpretation of producing better AUC score than G4Detector (section “Performance of G4-Attention on G4 Propensity Prediction”, which is present in Fig. 10 (c). similar to random and dishuffle, for PQ importance of inner Gs compared to outer Gs is higher as mentioned in Lee *et al*. [81]. Attribution scores for three negative sets (random, dishuffle and PQ) of G4-Attention trained using the *K*^+^ + PDS stabilizer dataset are depicted in S1 Fig (a), (b), and (c), which is present in the supplementary information.

The mutation maps reveal that G4-Attention is highly sensitive to mutations in inner G-rich regions separated by short loops, as shown in Figs. 10 (a), (b), and (c) respectively. This result is in agreement with IG methods, which gave high positive attribution scores to these inner G-rich regions. Moreover, in the same regions, mutating to G in positions 37, 44, 45, or 50 which connect G-tracts, increases the predicted probability of the G4-Attention model trained on random negatives, although mutating the flanking areas, has hardly any effect on the prediction [59]. These results indicate that the model trained on random negatives identifies regions of high guanine concentration, as opposed to the models trained on dishuffle and PQ negative sets, which exhibit sensitivity to nucleotides in the flanks [59]. The mutation map for three negative sets (random, dishuffle, and PQ) of G4-Attention trained using the *K*^+^ + PDS stabilizer dataset is depicted in S1 Fig (a), (b), and (c), which is present in the supplementary information.

### G4-folding principals learned by G4-Attention

In this section, we address the second part of our RQ-4. Here, we follow the work Barshai *et al*. [65] to conduct the necessary experiments to interpret G4-Attention. We examined how loop lengths, mismatches within the G-tracts, and the nucleotide composition of the flanking sequence influenced the prediction of mismatch scores.

#### Effect of Loop Lengths

We varied the length of the loops of the sequence of shape GGGNGGGNGGGNGGG from 1nt to 12nt for the G4-Attention trained on 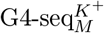 dataset is depicted in Fig. 11 (a). In Fig. 11 (a), loop1 and loop3 denote two outer loops and loop2 denotes as inner loop. We clearly observe that the G4-Attention 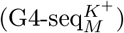 model predicts a reduction in mismatch score as the loop length increases, with a more pronounced effect in the inner loop (loop 2 in Fig. 11 (a)) in comparison to two outer loops. The opposite relationship between G4 stability and loop length, which G4-Attention learned from G4-seq data, aligns with findings from a previous low-throughput assay [71]. The effect of loop lengths using G4-Attention 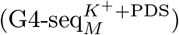 model is shown in S2 Fig (a), present in the supplementary information.

**Fig 11.**
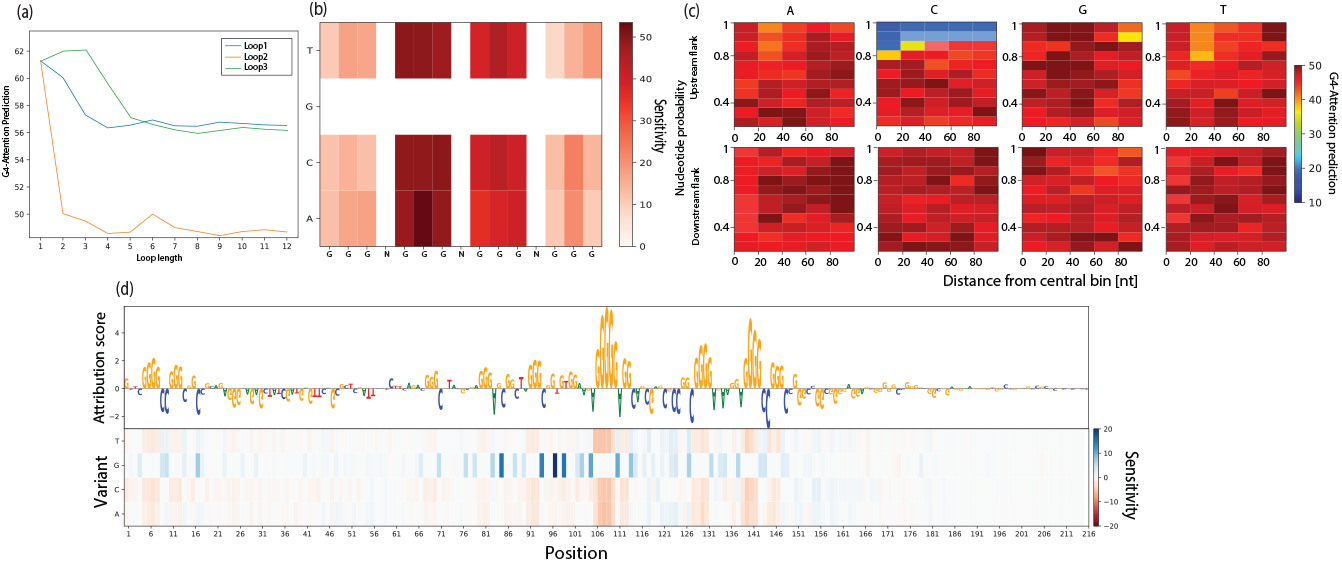
Interpreting G4-Attention. (a) Exploring the impact of loop length on predicting mismatch scores, we adjust the length of each loop within the canonical G4 sequence GGGNGGGNGGGNGGG individually and use G4-Attention, which has been trained on 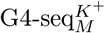, to predict the mismatch scores. (b) Impact of mismatch in G-tracts analyzed through G4-Attention predictions, which are based on 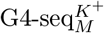. In this study, we focus on a standard G4 sequence, GGGNGGGNGGGNGGG, and individually mutate each nucleotide within the G-tracts to every other nucleotide, aiming to determine the variation in the predicted mismatch score. (c) Nucleotide composition effect in the flanking sequences on G4-Attention predictions, which is trained on 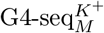. We take the canonical G4 of the form GGGNGGGNGGGNGGG and varied the probability of each nucleotide at a time, while assigning uniform probabilities to the other three nucleotide, in 20nt-long regions away from the central bin. We predict the mismatch score for each such variant, both for upstream and downstream flanks separately. (d) The mutation map shows the sensitivity of the G4-Attention trained on 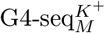 to mutations and the corresponding attributions report the importance of a given features of the model’s prediction.

#### Effect of Mismatch in the G-tracts

Here, we mutated G-tracts of GGGNGGGNGGG-NGGG for G4-Attention 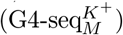 which is shown in Fig. 11 (b), which led to a reduction in the predicted mismatch score. Notably, the most significant decrement was observed for mutations in the second G-tract, compared to the canonical G4, which was initially observed in Barshai *et al*. [65] also shown by our model G4-Attention. We anticipated that mutations in the middle G-quartet would have a greater impact on G4 destabilization, as previously demonstrated in a low-throughput experiment [82]. We speculate that the above theory was not observed in our visualization of G4-Attention-learnt G4 folding principles as a consequence of a technological artifact of the G4-seq assay, which should be further studied for through mitigation, which also present in G4mismatch [65]. Effect of Mismatch in the G-tracts using G4-Attention 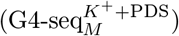 model is shown in S2 Fig (b), present in the supplementary information.

#### Effect of Nucleotide Composition

Here, we evaluated the impact of nucleotide composition in the flanking sequences of a 15nt-long motif, GGGNGGGNGGGNGGG, on mismatch score predictions using G4-Attention 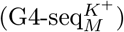, as illustrated in Fig. 11 (c). We observed that G4-Attention was more sensitive to changes in the upstream flanking sequence compared to the downstream flanking sequence. Specifically, increasing the probability of cytosine significantly reduced the sequence’s ability to form a G4, whereas increasing the probability of guanine had a lesser impact. For example, setting the cytosine probability to 1 and all other base pair probabilities to 0 within a 20nt upstream region of the central 15nt motif (GGGNGGGNGGGNGGG) resulted in a 60.29% decrease in the predicted mismatch score. Conversely, setting the guanine probability to 1 in the same region led to only a 5.32% increase in the predicted mismatch score. We propose that once a potential G4 structure reaches a certain level of stability, adding more guanines to an already stable configuration, like a canonical G4, has a limited impact on its stability. On the other hand, introducing cytosines increases the probability of guanines to form base pairs. This type of result was previously observed in the work by Barshai *et al*. [65]. Adenine and thymine exhibited similar effects and had a neutral effect on G4-Attention’s predictions. Modifying the probability of adenine and thymine in the region closest to the central 15nt-long i.e., in the range of 20nt upstream from the central 15nt-long canonical G4 bin, increases the predicted mismatch score. A possible reason could be the fact that increasing their probability corresponds to a decrease in cytosine probability, this was previously observed by G4mismatch [65]. Effect of Nucleotide Composition using G4-Attention 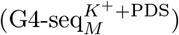 model is shown in S2 Fig (c), present in the supplementary information.

#### Key Features Identification of G4-Attention of a Genome Sequence

Finally, we employed the integrated gradient (IG) method to visualize the key features recognized by G4-Attention 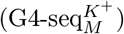. Fig. 11 (d) shows the attribution score calculated for the KRAS proto-oncogene using G4-Attention trained on *K*^+^ stabilizer. We take KRAS proto-oncogene because in the previous work G4mismatch [65] this particular genome sequence is used to conduct the current study. The attribution score indicates that G4-Attention prediction is based on G-rich subsequence. The relevant guanines reside in continuous stretches of tall letters. Again, the corresponding mutation map supports the results obtained by the IG method, showing that the model is highly sensitive to mutation of a given nucleotide from or to guanine, which is also clearly present in the work G4mismatch [65]. Attribution score and mutation map of KRAS proto-oncogene using G4-Attention 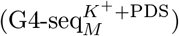 model are shown in S2 Fig, present in the supplementary information.

### Ablation Study

Here, we perform a series of ablation experiments on class imbalanced dataset G4-seq_IB_ for our proposed model G4-Attention. Initially, we consider CNN as our baseline model. Here, we take 1D CNN to extract features from the genome sequence and then we predict whether the genome sequence has G4 propensities or not. It achieves an AUPRC score of 0.699. Subsequently, we introduce an LSTM layer on top of the CNN layer and this model produces an AUPRC score of 0.742. Furthermore, we introduce an attention layer on the CNN + LSTM model, which increases the model performance as it produces an AUPRC score of 0.759. Finally, we implement our model G4-Attention by replacing the LSTM layer in the CNN + LSTM + Attention model with a Bi-LSTM layer and this architecture produces an AUPRC score of 0.771. Fig. 12 shows that G4-Attention contributes significantly towards achieving state-of-the-art results on this task. Here, we take G4-seq_IB_ for conducting ablation experiments because the prediction of positive G4s on the datasets with high negative skewness is a challenging task. Since the dataset is class imbalanced, we take the AUPRC metric to conduct the ablation experiments.

**Fig 12.**
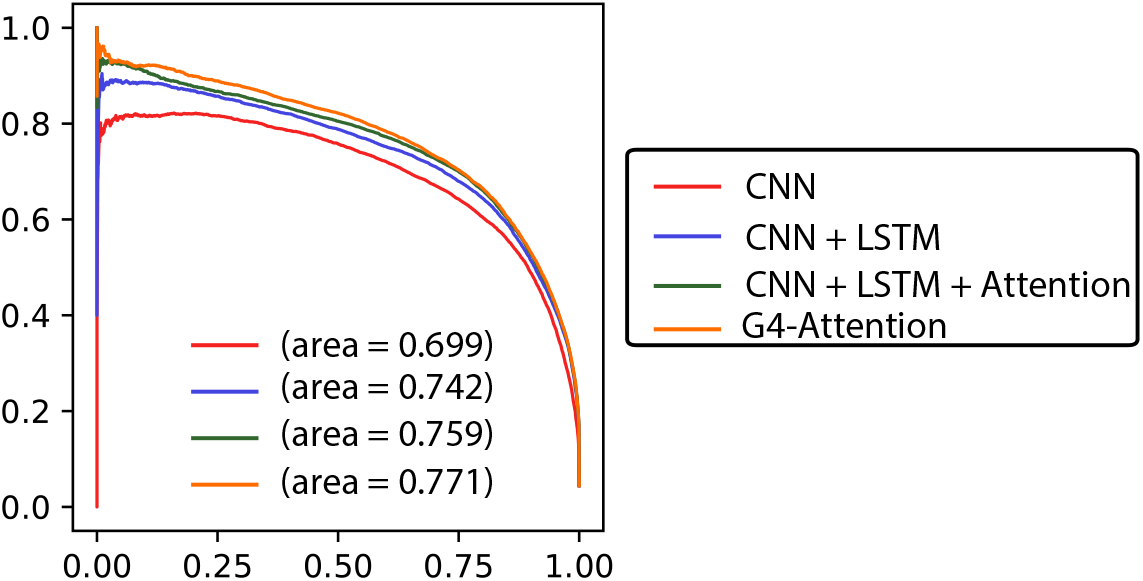
A series of ablation studies were performed on the class imbalance dataset G4-seq_IB_. The precision-recall curves (AUPRC) have been shown.

## Conclusion

In this work, we present a novel framework G4-Attention for the task of predicting G4 formation in a genome sequence. Our model uses CNN, Bi-LSTM, and an attention-based network to predict the G4 formation of a genome sequence. To investigate our model performance, we carried out different kinds of experiments such as G4-prediction task and G4-mismatch score prediction task on various kinds of datasets including balanced datasets, regression datasets, class imbalanced datasets with high negative skewness, and cell-type specific datasets. Various kinds of experimental results analysis show that our model achieves new state-of-the-art results on all these datasets and a significant prediction improvement has also been observed with respect to other available benchmark models in the literature. Notably, besides the *in vitro* prediction, G4-Attention can also predict the G4s very accurately in cell-type specific DNA sequences. Next, to test the performance of our model in out-domain scenarios, we extended our study to the different kinds of non-human genomic sequences of species like mouse, zebrafish, drosophila, and others. Experimental results show that our model trained on the human genome not only produces excellent results on in-domain test data but also produces state-of-the-art results for out-domain test data, suggesting the potential use of G4-Attention on diverse species in the context of G4s prediction. All these observations collectively indicate that we have successfully developed a deep-learning model, G4-Attention with Bi-LSTM and attention layers using human genome sequences which can predict the G4s much more accurately in comparison to other available benchmark models.

## Supporting information

**S1 Table**. Species name, their scientific name and the number of instances present for each of the three negative sets, for each of the species present in G4-seq_B_ for *K*^+^ and *K*^+^ + PDS used in this study.

**S2 Table**. Species name, their scientific name and the number of G4 mismatch scores reported for each of the species present in 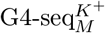 and 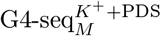 used in this study.

**S1 Fig**. Example of attribution scores assigned by the integrated gradient method and mutation maps for one given sequence based on models trained on the G4-seq_B_ dataset for one genome sequence. Figures (a), (b) and (c) shows that G-stretches, separated by short loops, receive relatively high scores for models trained on the Random, Dishuffle and PQ negative sets.

**S2 Fig**. Interpreting G4-Attention. (a) Effect of loop length on mismatch score prediction. Here we take a canonical G4 GGGNGGGNGGNGGG and varied the length of each loop seprately, and predict the mismatch score using G4-Attention trained on 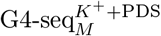. (b) Effect of mismatch in the G-tracts using G4-Attention predictions trained on 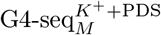. Here we take the same canonical G4 which is GGGNGGGNGGNGGG, and mutated each nucleotide in the G-tracts separetly to all other nucleotide, and calculate the change in the predicted mismatch score. (c) Nucleotide composition effect in the flanking sequences on G4-Attention predictions, which is trained on 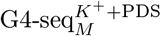. We take the canonical G4 of the form GGGNGGGNGGNGGG and varied the probability of each nucleotide at a time, while assigning uniform probabilities to the other three nucleotide, in 20nt-long regions away from the central bin. We predict the mismatch score for each such variant, both for upstream and downstream flanks separately. (d) The mutation map shows the sensitivity of the G4-Attention trained on 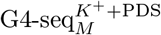 to mutations and the corresponding attributions report the importance of a given features of the model’s prediction.

## Acknowledgments

We thank Prof. Yaron Orenstein for helping us to conduct the necessary experiments for the interpretation study and for generating negative sets for cross-domain datasets. We also thank Minhazuddin Ahmed for helping us to generate images for the interpretation study. This study was funded by the Indian Association for the Cultivation of Science (IACS), Kolkata, India. SM thanks IACS for a research fellowship. PP thanks to the Council of Scientific and Industrial Research (CSIR), India for a Senior Research Fellowship.

## Footnotes

1 Mouse, zebrafish, drosophila, etc. are used in the case study.

2 https://cran.r-project.org/web/packages/gkmSVM.

3 http://genome.ucsc.edu/

4 As mentioned in this work [61]

5 https://pytorch.org/ignite/index.html

6 Here we use G4NN, as in the original paper by Korsakova et. al [61] used the model without epigenetic features i.e; the vanilla version as G4NN.

