## Supplementary material for "G4-Attention: Deep Learning Model with Attention for predicting DNA G-Quadruplexes": S1 Figure, S2 Figure, S1 Table, S2 Table

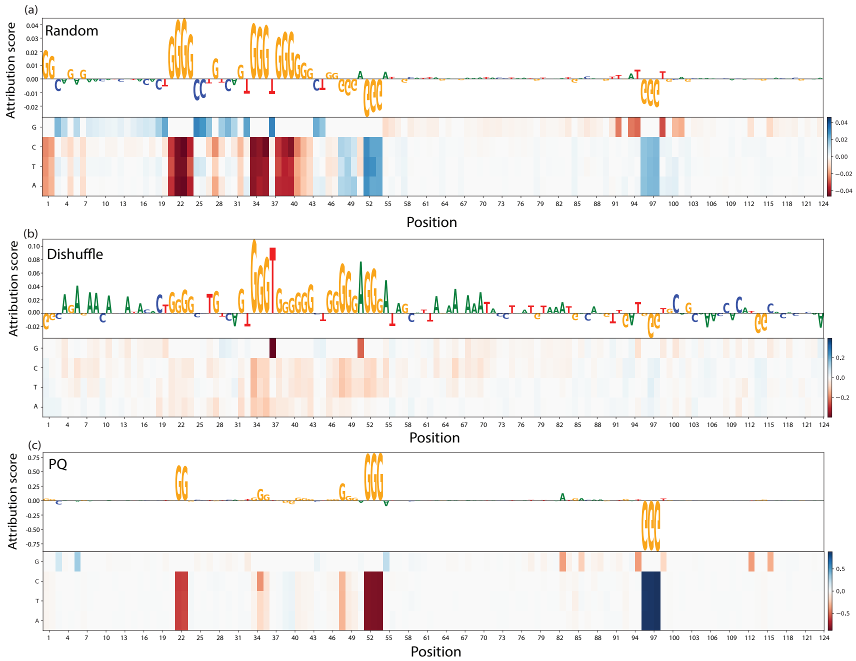


**S1 Fig.** Example of attribution scores assigned by the integrated gradient method and mutation maps for one given sequence based on models trained on the G4-seq$_B$ dataset for one genome sequence. Figures (a), (b) and (c) shows that G-stretches, separated by short loops, receive relatively high scores for models trained on the Random, Dishuffle and PQ negative sets.

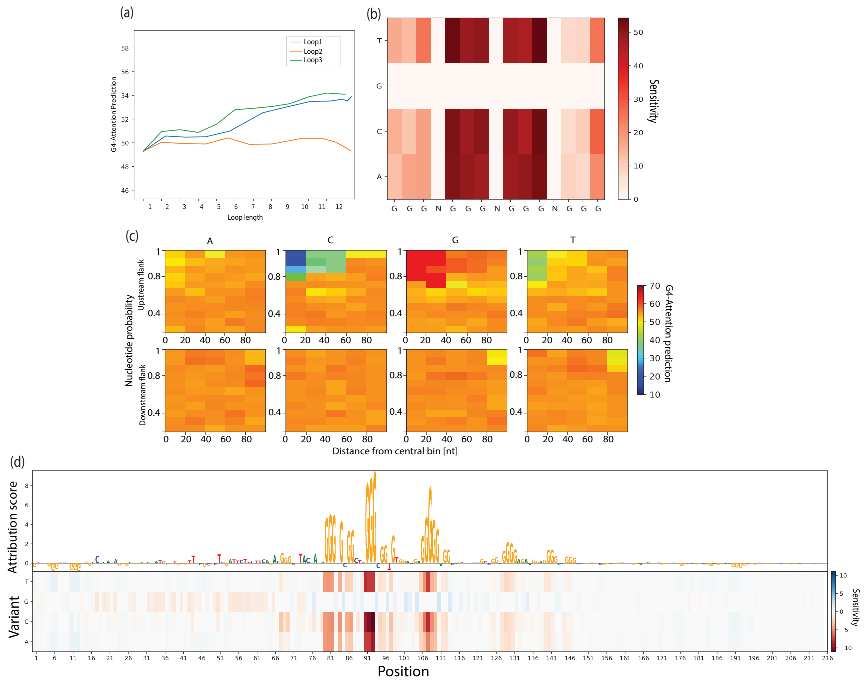


**S2 Fig.** Interpreting G4-Attention. (a) Effect of loop length on mismatch score prediction. Here we take a canonical G4 GGGNGGGNGGNGGG and varied the length of each loop seprately, and predict the mismatch score using G4-Attention trained on G4-seq$_M^{K^+ + \mathrm{PDS}}$. (b) Effect of mismatch in the G-tracts using G4-Attention predictions trained on G4-seq$_M^{K^+ + \mathrm{PDS}}$. Here we take the same canonical G4 which is GGGNGGGNGGNGGG, and mutated each nucleotide in the G-tracts separetly to all other nucleotide, and calculate the change in the predicted mismatch score. (c) Nucleotide composition effect in the flanking sequences on G4-Attention predictions, which is trained on G4-seq$_M^{K^+ + \mathrm{PDS}}$. We take the canonical G4 of the form GGGNGGGNGGNGGG and varied the probability of each nucleotide at a time, while assigning uniform probabilities to the other three nucleotide, in 20nt-long regions away from the central bin. We predict the mismatch score for each such variant, both for upstream and downstream flanks separately. (d) The mutation map shows the sensitivity of the G4-Attention trained on G4-seq$_M^{K^+ + \mathrm{PDS}}$ to mutations and the corresponding attributions report the importance of a given features of the model's prediction.

**S1 Table.** Species name, their scientific name and the number of instances present for each of the three negative sets, for each of the species present in G4-$seq_B$ for $K^+$ and $K^+$ + PDS used in this study.

| **Species** | **Scientific Name** | **Negative Type** | **G4-$seq_B$ ($K^+$)** | **G4-$seq_B$ ($K^+$ + PDS)** |
|---|---|---|---|---|
| Human | *Homo sapiens* | Random | 838,992 | 2,658,186 |
| | (*H. sapiens*) | Dishuffle | 868,516 | 2,752,808 |
| | | PQ | 860,033 | 1,491,271 |
| Mouse | *Mus musculus* | Random | 1,572,759 | 3,443,964 |
| | (*M. musculus*) | Dishuffle | 1,595,442 | 3,493,538 |
| | | PQ | 999,058 | 1,948,106 |
| Zebrafish | *Danio reiro* | Random | 281,800 | 640,925 |
| | (*D. reiro*) | Dishuffle | 281,208 | 641,488 |
| | | PQ | 165,303 | 345,443 |
| Drosophila | *Drosophila melanogaster* | Random | 38,720 | 110,325 |
| | (*D. melanogaster*) | Dishuffle | 38,794 | 110,514 |
| | | PQ | 31,483 | 67,343 |
| Roundworm | *Caenorhabditiselegans* | Random | 8,288 | 21,552 |
| | (*C. elegans*) | Dishuffle | 8,288 | 21,552 |
| | | PQ | 4,144 | 10,776 |
| Brewer's yeast | *Saccharomyces cerevisiae* | Random | 206 | 11,278 |
| | (*S. cerevisiae*) | Dishuffle | 206 | 11,278 |
| | | PQ | 103 | 10,776 |
| Leishmania | *Leishmania major* | Random | 34,686 | 73,882 |
| | (*L. major*) | Dishuffle | 34,686 | 73,882 |
| | | PQ | 17,343 | 36,941 |
| Plasmodium | *Plasmodium falciparum* | Random | 652 | 346 |
| | (*P. falciparum*) | Dishuffle | 652 | 346 |
| | | PQ | 326 | 173 |
| Thale cress Plant | *Arabidopsis thaliana* | Random | 4,814 | 23,906 |
| | (*A. thaliana*) | Dishuffle | 4,814 | 23,906 |
| | | PQ | 2,407 | 11,953 |
| E. coli | *Escherichia coli* | Random | 94 | 1,120 |
| | (*E. coli*) | Dishuffle | 94 | 1,120 |
| | | PQ | 47 | 560 |
| Rhodobacter | *Rhodobacter sphaeroides* | Random | 4,582 | 582 |
| | (*R. sphaeroides*) | Dishuffle | 4,582 | 582 |
| | | PQ | 2,291 | 291 |
| Trypanosoma | *Trypanosoma brucei* | Random | 6,472 | 21,332 |
| | (*T. brucei*) | Dishuffle | 6,472 | 21,332 |
| | | PQ | 3,236 | 10,666 |

**S2 Table.** Species name, their scientific name and the number of G4 mismatch scores reported for each of the species present in G4-seq$_M^{K^+}$ and G4-seq$_M^{K^+ + \mathrm{PDS}}$ used in this study

| **Species** | **Scientific Name** | **G4-seq$_M^{K^+}$** | **G4-seq$_M^{K^+ + \mathrm{PDS}}$** |
|---|---|---|---|
| Human | *Homo sapiens* (*H. sapiens*) | 372,627,364 | 377,015,360 |
| Mouse | *Mus musculus* (*M. musculus*) | 315,325,949 | 316,404,581 |
| Zebrafish | *Danio reiro* (*D. reiro*) | 165,668,249 | 152,358,421 |
| Drosophila | *Drosophila melanogaster* (*D. melanogaster*) | 15,870,402 | 15,899,380 |
| Roundworm | *Caenorhabditiselegans* (*C. elegans*) | 13,365,121 | 13,160,158 |
| Brewer's yeast | *Saccharomyces cerevisiae* (*S. cerevisiae*) | 1,589,250 | 1,551,955 |
| Leishmania | *Leishmania major* (*L. major*) | 4,338,042 | 4,342,208 |
| Plasmodium | *Plasmodium falciparum* (*P. falciparum*) | 1,828,562 | 1,657,136 |
| Thale cress Plant | *Arabidopsis thaliana* (*A. thaliana*) | 15,332,638 | 15,335,473 |
| E. coli | *Escherichia coli* (*E. coli*) | 603,254 | 603,299 |
| Rhodobacter | *Rhodobacter sphaeroides* (*R. sphaeroides*) | 597,896 | 600,911 |
| Trypanosoma | *Trypanosoma brucei* (*T. brucei*) | 2,731,786 | 2,735,227 |
